# Lipid-Protein Reciprocal Coupling: A Quantitative Analysis of Membrane Curvature, Architecture, and Protein Association

**DOI:** 10.64898/2026.09.24.751570

**Authors:** Md Sorique Aziz Momin, Debanjan Sarkar, Nayan De, Debmalya Bhunia, Achinta Sannigrahi

## Abstract

Cellular membranes are dynamic molecular landscapes in which lipid composition, transbilayer asymmetry, membrane curvature, thickness and mechanics collectively shape protein organization and function, while proteins can in turn remodel lipid organization and membrane architecture. Yet lipid-protein interactions are often treated as discrete binding events, obscuring the extent to which the physical organization of the membrane itself constitutes an active determinant of protein conformational states and cellular activity. Here, we investigate lipid-protein reciprocal coupling by integrating quantitative analyses of curvature-dependent protein-membrane association and hydrophobic-run organization with established structural, lipidomic and biochemical evidence. Our analysis reveals substantial variation in protein association with curved versus flat membrane environments and highlights how membrane curvature, together with lipid composition, cholesterol and asymmetry, can regulate protein structure and function, including in disease-associated systems. We further examine emerging lipid sensors, spatial proteomics and advanced biomimetic membrane platforms that are beginning to bridge molecular mechanisms with protein-lipid organization in cells. In particular, suspended membranes, pore-spanning systems and nanopillar architectures offer opportunities to independently control lipid composition, leaflet asymmetry, curvature and mechanics while directly interrogating protein behaviour. Together, these findings support a model in which membrane architecture encodes physical information that is read by proteins and reciprocally reshaped by protein activity to regulate cellular function.

## Introduction

Cellular membranes are dynamic lipid environments that continuously change and shape cellular organization and function. Their composition, spatial organization and mechanics collectively shape membrane-protein function and cell signaling. Rather than serving as a passive solvent for membrane proteins, the lipid bilayer provides a chemically and physically heterogeneous environment defined by lipid identity, acyl-chain composition, membrane thickness, curvature, electrostatics, asymmetry and mechanical properties. These features vary substantially between organelles and can change locally and dynamically within a membrane. Consequently, membrane proteins experience a lipid environment fundamentally different from the relatively homogeneous aqueous environment surrounding soluble proteins, and their structure and function are shaped by this local membrane context ^1,2^. Early studies of hydrophobic mismatch established that differences between the hydrophobic length of a transmembrane protein and the thickness of the surrounding bilayer can alter protein orientation, oligomerization and conformational dynamics, while simultaneously reorganizing lipid packing and membrane thickness ^3,4^. Specific lipids can additionally act as molecular regulators. Phosphoinositides, for example, bind defined sites on ion channels and stabilize functional conformations, whereas cholesterol and anionic phospholipids can modulate membrane-protein structure and activity through both specific and collective interactions ^2,5^.

However, lipid regulation of proteins represents only one direction of the interaction. Membrane proteins also actively reshape the lipid environment that surrounds them. Protein insertion, oligomerization and conformational changes can perturb local lipid packing, membrane thickness, curvature and lipid distribution. The resulting lipid organization can, in turn, influence the behavior of the same or neighboring proteins. This reciprocal relationship was recognized in early biophysical models of lipid-protein interactions and has since been supported by experimental and computational studies showing that proteins can alter the organization, dynamics and phase behavior of surrounding lipids^4^. Membrane curvature provides a particularly clear example of this reciprocity. Lipid composition and packing determine the energetic cost of membrane bending, whereas proteins can sense, stabilize or generate curvature through amphipathic helices, scaffolding domains and oligomerization. BAR-domain proteins exemplify this principle, in which membrane geometry influences protein recruitment while protein assembly can subsequently stabilize or generate membrane curvature^6^. These findings motivate lipid-protein reciprocal coupling as a framework in which lipid-protein interactions extend beyond discrete binding events to encompass the coupled chemical and physical organization of membranes. Such coupling operates across spatial scales, from specific lipid contacts at individual binding sites to nanoscale lipid shells and larger-scale changes in membrane curvature, domains and organization. Recent advances in cryo-electron microscopy, native mass spectrometry, lipidomics, super-resolution imaging and molecular simulations have substantially expanded our ability to interrogate lipid-protein interactions. Yet capturing their dynamic and real time reciprocal nature remains challenging. Structural and biochemical approaches can identify specific lipid contacts but often provide static or ensemble-averaged views, whereas cellular approaches preserve membrane complexity but make it difficult to distinguish direct lipid interactions from indirect effects of local membrane organization. Moreover, lipid composition, leaflet asymmetry, curvature, packing and mechanics are tightly coupled, making it difficult to isolate the contribution of individual membrane properties. For example, combining native mass spectrometry with lipidomics can identify which lipids remain associated with membrane proteins and reveal preferences for specific lipid headgroups and acyl chains ^7^. However, these methods provide limited information about how such interactions change within a dynamic membrane.

Understanding how lipid identity and membrane physical properties influence protein behavior, and how proteins in turn remodel their surrounding lipids, therefore requires approaches that combine molecular specificity, spatial resolution and control over membrane architecture.

This challenge is particularly important because conventional reductionist approaches often require detergent solubilization or reconstitution into simplified symmetric bilayers, potentially removing the asymmetry, curvature, lipid heterogeneity and mechanical stresses that regulate protein behavior. Conversely, measurements in living cells preserve physiological complexity but frequently lack sufficient molecular resolution to distinguish direct lipid contacts from indirect effects of membrane organization. Bridging these scales therefore requires complementary approaches that combine lipid-specific probes, spatial proteomics, molecular simulations and advanced imaging with biomimetic systems in which lipid composition, leaflet asymmetry, curvature, membrane thickness and mechanics can be systematically controlled. Such strategies offer a route toward connecting molecular interactions with the emergent properties of cellular membranes.

Here, we investigate lipid-protein reciprocal coupling by combining a quantitative analysis of curvature-dependent protein-membrane association with published structural, biochemical and lipidomic evidence. We first establish the physicochemical framework linking lipid diversity, asymmetry, cholesterol, membrane curvature, and hydrophobic matching. We then quantitatively compare curved- and flat-membrane association across a structurally diverse set of membrane proteins and examine how these trends relate to known mechanisms of membrane remodeling and protein function. Finally, we integrate these findings with disease-associated protein transitions and emerging experimental platforms capable of directly testing reciprocal lipid-protein coupling.

### Membrane lipids evolved to create structural-functional landscapes

Membrane lipids evolved to create functional landscapes that organize proteins, regulate their activity, and enable cells to sense and respond to their environment. These are essential in all domains of life because they separate the cell from its surroundings, maintain chemical gradients and provide an organized platform for transport, metabolism and signalling. Although these basic functions are shared, membrane lipid composition differs across archaea, bacteria and eukaryotes, reflecting their distinct evolutionary histories and physiological needs. Phylogenomic analyses suggest that some phospholipid-biosynthetic enzymes, including CDP-alcohol transferases, may have been present in the last universal common ancestor, or cenancestor^8^. During the subsequent divergence of life, archaeal and bacterial membranes became chemically distinct. **Figure 1A** shows the distinct phospholipid biosynthetic routes used by the major domains of life, with archaeal membranes formed mainly from *sn*-glycerol-1-phosphate linked by ether bonds to methyl-branched isoprenoid chains and bacterial and eukaryotic membranes based largely on *sn*-glycerol-3-phosphate ester-linked to linear fatty acids ^8^. Hydrothermal environments may have further supported Fischer-Tropsch-type reactions in which carbon monoxide and hydrogen formed hydrocarbons in the presence of mineral catalysts, followed by the production of long-chain fatty acids and alcohols that contributed to primitive membrane formation ^9–12^. Several models have been proposed to explain lipid evolution, including independent evolution from an acellular cenancestor, evolution within mineral-bounded compartments, specialization from pre-cells with heterochiral membranes and selection-pressure driven lipid synthesis from an ancestor with an achiral membrane (**Figure 1B**)^8,13–15^. **Figure 1B** summarizes these four proposed models for the origin and divergence of archaeal and bacterial membrane lipids. Early membranes may also have formed from simple amphiphilic molecules, such as fatty acids and alcohols, rather than from the complex phospholipids found in modern cells. These molecules could have been supplied by carbonaceous meteorites or generated through prebiotic chemistry and could spontaneously assemble into vesicles capable of encapsulating biomolecules, maintaining chemical gradients, growing and dividing. Following the endosymbiotic origin of mitochondria, eukaryotic membranes evolved with increased complexity through the emergence of specialized organelles and diverse lipid compositions. This compartmentalization enabled distinct membrane functions, supporting advanced cellular processes such as trafficking, signaling, and energy metabolism. More broadly, membrane lipid composition has expanded from the relatively simple phospholipid-rich membranes of bacteria and host-derived envelopes of viruses to the highly diverse glycerophospholipids, sphingolipids, glycolipids and sterols of animal cells^1,16–18^. Collectively, these evolutionary models illustrate how lipid composition became adapted to distinct cellular and environmental requirements. The homeoviscous adaptation model further explains how organisms maintain optimal membrane function by adjusting lipid composition to preserve membrane fluidity under changing environmental conditions (**Figure 1C**). By altering the ratio of saturated and unsaturated fatty acids, chain length, and lipid headgroup composition, cells regulate membrane viscosity and ensure proper function of membrane-associated proteins and processes^19^. In parallel to this homeoviscous adaptation model, deep-sea ctenophores and other invertebrates alter the curvature properties of their membrane phospholipids to withstand high pressure and low temperature, enriching plasmalogens, ether phospholipids and long, highly unsaturated acyl chains while reducing lysolipids^20^. This process, termed homeocurvature adaptation, helps preserve membrane organization under extreme conditions (**Figure 1D**), whereas rapid transfer to surface pressure can disrupt membrane integrity through lipid phase transitions^20^. These recent findings point to a close relationship between lipid biosynthesis, genomic variation, membrane composition, leaflet asymmetry and curvature, which together shape membrane organization and cellular function. These evolutionary and adaptive changes provide the basis for examining how plasma-membrane asymmetry and bilayer geometry emerge from lipid diversity and influence membrane morphology, membrane-protein function and cell signalling. Lipid diversity did not merely increase membrane complexity; it created distinct physical environments that can be exploited by membrane proteins. In modern eukaryotic membranes, this lipid diversity is further organized spatially, not only among cellular organelles but also between the two leaflets of individual bilayers. This transbilayer organization converts differences in lipid chemistry into differences in packing, fluidity, thickness and curvature, providing the physical basis for the membrane asymmetry and curvature-dependent protein association analyzed in the subsequent sections.

**Figure 1.**
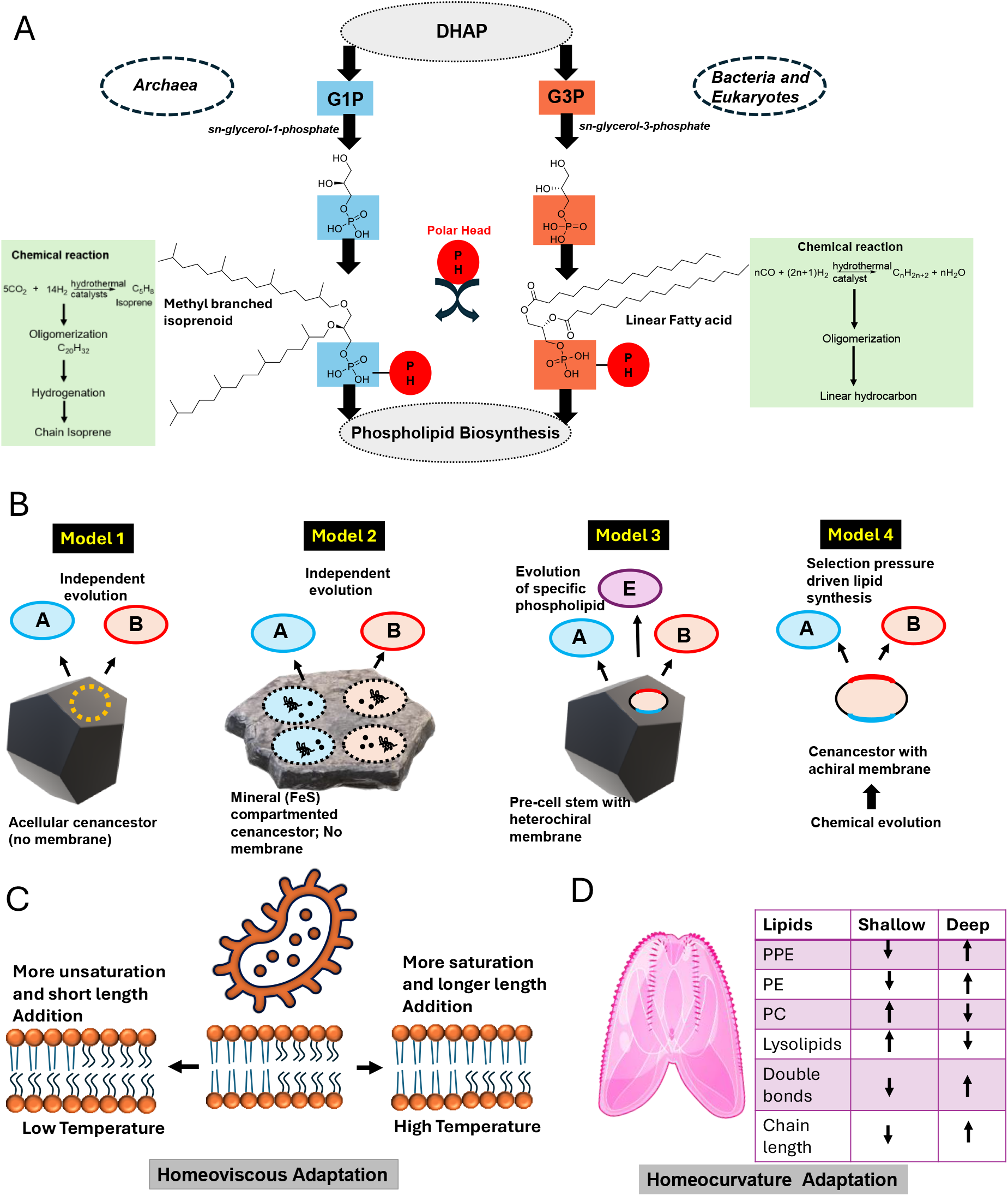
Lipid evolutionary divergence, homeoviscous, and homeocurvature adaptation mechanisms. **(A)** Phospholipid biosynthetic pathways originating from a central dihydroxyacetone phosphate (DHAP) precursor. In *Archaea*, conversion to sn-glycerol-1-phosphate (G1P) and linkage with methyl-branched isoprenoids via hydrothermal catalysts forms ether-linked membrane boundaries. In *Bacteria and Eukaryotes*, conversion to sn-glycerol-3-phosphate (G3P) and esterification with linear fatty acids generated from chemical evolution pathways produces standard ester-linked membrane bilayers containing distinct polar heads (P H). **(B)** Evolutionary models depicting membrane development from a common ancestor to separate cellular domains, illustrating an acellular cenancestor with no true membrane (Model 1), a mineral (FeS) compartmented cenancestor (Model 2), a pre-cell stem organism containing a heterochiral mixed membrane (Model 3), and a selection pressure-driven lipid pathway originating from an achiral membrane progenitor (Model 4). **(C)** Homeoviscous adaptation pathway of cellular membranes adjusting to fluctuating thermal environments. Cold conditions promote shorter chain lengths and high unsaturation addition to maintain fluidity, whereas hot environments promote longer, highly saturated additions to stabilize the bilayer. **(D)** Homeocurvature adaptation profile tracking structural alterations in deep-sea environments to accommodate hydrostatic stress. The matrix details the shifting concentrations of inverse-cone lipids (PPE, PE), cylindrical lipids (PC, lysolipids), and tail configurations (double bonds, chain length) required to maintain structural integrity from shallow to deep zones.

### Lipid asymmetry creates a physical membrane landscape

One major consequence of this evolved lipid diversity is the asymmetric organization of lipids across the two leaflets of the plasma membrane, which creates distinct physical environments for membrane-associated proteins. The plasma membrane (PM) of mammalian cells displays a remarkable transbilayer lipid asymmetry that is actively established by lipid biosynthesis and maintained by ATP-dependent lipid transporters, including P4-ATPase flippases, ABC transporters (floppases), and Ca²⁺-activated scramblases^21–23^. Early phospholipase digestion studies using intact erythrocytes first demonstrated that phosphatidylcholine (PC) and sphingomyelin (SM) are almost exclusively localized to the exoplasmic leaflet, whereas phosphatidylethanolamine (PE) and phosphatidylserine (PS) are concentrated in the cytoplasmic leaflet ^24,25^. Subsequent quantitative analyses combining membrane-impermeable phospholipase A₂, phospholipase C and sphingomyelinase digestion, which selectively hydrolyze outer leaflet phospholipids, with chemical derivatization using trinitrobenzene sulfonate (TNBS) or sulfo-NHS reagents, selective extraction, and quantitative LC-MS/MS lipidomics to identify the remaining inner leaflet lipids have shown that approximately 75-85% of total PC and >90% of SM reside in the outer leaflet, whereas 80-90% of PE and >95% of PS are restricted to the inner leaflet(**Figure 2A**). Phosphatidylinositol (PI) and its phosphorylated derivatives, including PI(4)P and PI(4,5)P₂, are found almost exclusively on the cytoplasmic leaflet, where they recruit numerous signaling and cytoskeletal proteins^26,27^. Complementary measurements using fluorescence quenching, environment-sensitive dyes, spin-label electron paramagnetic resonance (EPR), neutron scattering, and molecular dynamics simulations further demonstrated that the exoplasmic leaflet is more ordered, thicker and mechanically rigid, whereas the cytoplasmic leaflet is thinner, more disordered and considerably more dynamic ^27^.

**Figure 2.**
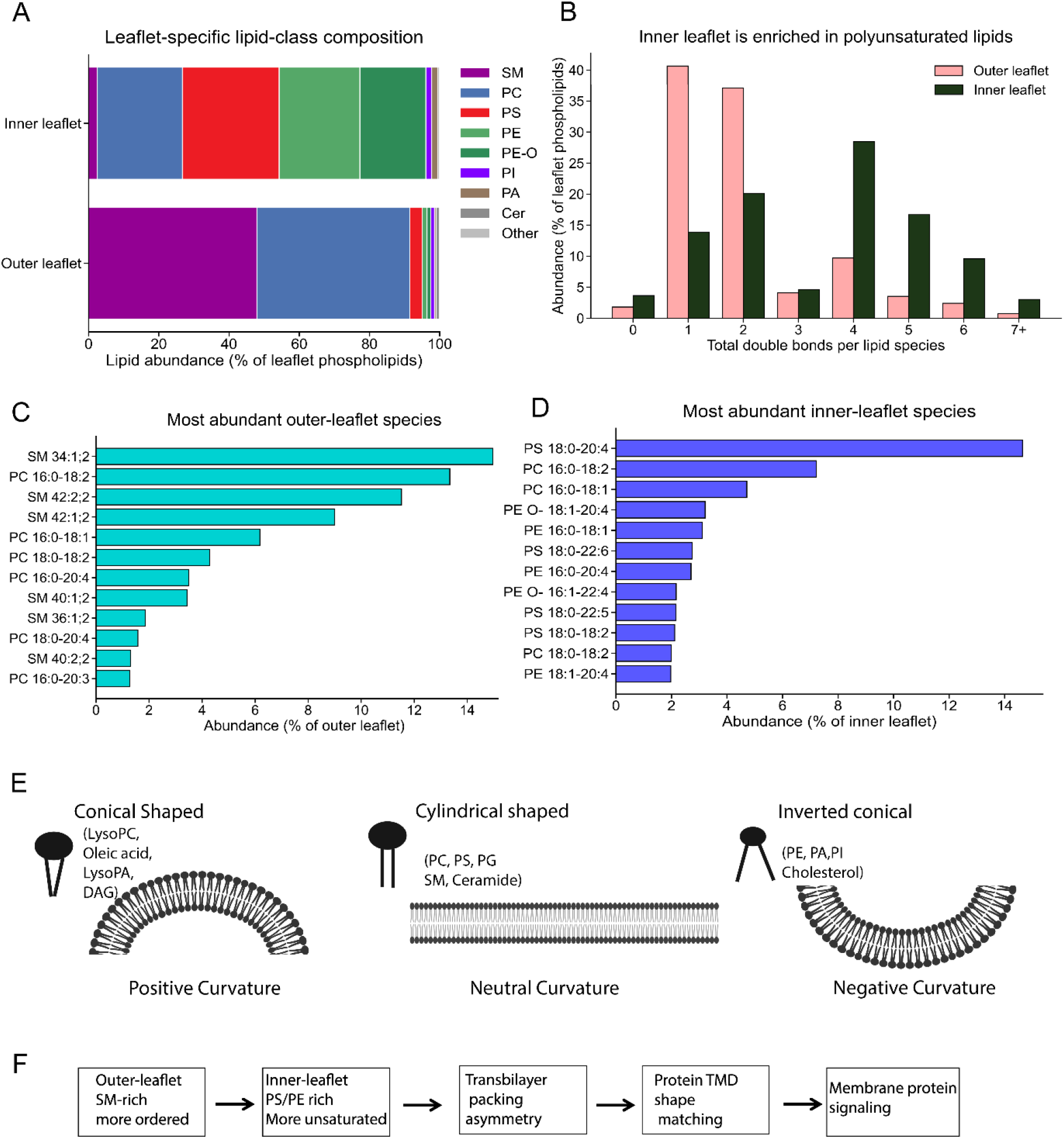
Leaflet-specific lipid class distributions, geometric curvature mechanics, and asymmetry dynamics. **(A)** Leaflet-specific lipid-class composition illustrating enrichment of sphingomyelin (SM) and phosphatidylcholine (PC) in the outer leaflet and phosphatidylethanolamine (PE), phosphatidylserine (PS), and phosphatidylinositol (PI) in the inner leaflet. **(B)** Distribution of phospholipid unsaturation across the two leaflets, showing enrichment of polyunsaturated lipid species in the inner leaflet. **(C)** Most abundant lipid molecular species in the outer leaflet. **(D)** Most abundant lipid molecular species in the inner leaflet. Data shown in panels A-D were adapted from the plasma-membrane lipidomic analysis of Lorent et al^27^. **(E)** Schematic illustrating the relationship between lipid molecular geometry and preferred membrane curvature. Conical lipids, including LysoPC, oleic acid, LysoPA, and DAG, favor positive curvature; approximately cylindrical lipids, including PC, PS, PG, SM, and ceramide, favor relatively neutral curvature; and inverted-conical lipids, including PE, PA, PI, and cholesterol, favor negative curvature. **(F)** Conceptual model illustrating how differences in leaflet composition and lipid packing generate transbilayer structural asymmetry that can influence transmembrane-domain (TMD) shape matching and membrane-protein organization. The conceptual relationship between leaflet packing asymmetry and TMD structural asymmetry is based on findings reported by Lorent et al^27^.

Beyond phospholipid headgroups, the two membrane leaflets differ substantially in their acyl-chain composition. Comprehensive lipidomic analyses of human erythrocytes using selective enzymatic digestion followed by LC-MS/MS, revealed that the outer leaflet is enriched in saturated and monounsaturated phospholipids, particularly SM species such as d18:1/16:0, d18:1/24:0 and d18:1/24:1, together with PC species including PC 16:0/16:0, PC 16:0/18:1 and PC 18:0/18:1, resulting in a tightly packed, liquid-ordered environment ^27^. In contrast, the cytoplasmic leaflet contains a substantially higher proportion of polyunsaturated glycerophospholipids, including PE 18:0/20:4, PE 18:0/22:6, PS 18:0/20:4, PS 18:0/22:6, PI 18:0/20:4 and PI 18:0/22:6, which possess long-chain arachidonic (20:4) and docosahexaenoic (22:6) acyl chains that increase conformational flexibility and reduce lipid packing (**Figure 2B-D**). As a consequence, the average degree of unsaturation of the cytoplasmic leaflet is nearly two-fold higher than that of the exoplasmic leaflet, producing a membrane that is significantly more fluid despite the presence of similar cholesterol concentrations.

This structural asymmetry has profound functional consequences, regulating membrane curvature, vesicle budding, receptor signaling, mechano-transduction, cytoskeletal attachment and lipid-protein interactions. Moreover, disruption of lipid asymmetry through activation of scramblases results in the externalization of PS, a conserved signal for apoptosis, blood coagulation and immune recognition, highlighting that maintenance of lipid asymmetry is essential for plasma membrane integrity and cellular homeostasis ^23,26^.

Depending on this geometry, lipids can be broadly classified as cylindrical, cone-shaped, or inverted-cone-shaped molecules, each favoring distinct membrane curvatures. Cylindrical lipids, such as phosphatidylcholine and sphingomyelin, preferentially stabilize flat bilayers, whereas cone-shaped lipids including phosphatidylethanolamine favor negative curvature, and inverted-cone lipids such as lysophospholipids promote positive curvature (**Figure 2E**). These geometric preferences are commonly described by the critical packing parameter:

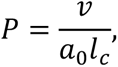

where *v* is the hydrocarbon-chain volume, *a*_0_ is the effective headgroup area, and *l_c_* is the hydrocarbon-chain length. Lipids with P ≈ 1 are generally cylindrical and favor lamellar bilayers, those with P < 1 tend to generate positive curvature, whereas lipids with P > 1 preferentially stabilize negatively curved membranes or non-lamellar structures ^28^. Although the critical packing parameter provides an intuitive molecular description of lipid geometry, the collective behavior of lipid mixtures is more appropriately described by spontaneous curvature (*C*₀), which reflects the preferred curvature of a membrane in the absence of external constraints. Differences in lipid composition between the two membrane leaflets therefore generate unequal spontaneous curvatures, producing transbilayer bending stresses that influence membrane remodeling and protein activity(**Figure 2F**).

The enrichment of saturated SM species in the outer leaflet favors tighter lipid packing and greater order, whereas the high abundance of polyunsaturated lipids in the inner leaflet produces a more loosely packed and fluid environment. At cellular length scales, membrane deformation is more accurately described by continuum elasticity theory. The energetic cost (*F*) of bending a membrane is commonly represented by the Canham-Helfrich free-energy equation:

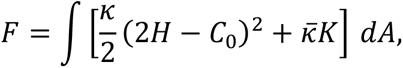

where *κ* is the bending rigidity, ***κ̅*** is the Gaussian curvature modulus, *H* and *K* are the mean and Gaussian curvatures, respectively, and *C*_0_ is the spontaneous curvature^29^.

Collectively, lipid composition, molecular geometry, leaflet asymmetry, and membrane mechanics define the physicochemical landscape experienced by membrane proteins. Variations in membrane thickness, curvature, lateral pressure, and lipid packing influence transmembrane-domain architecture, conformational dynamics, oligomerization, and protein partitioning into distinct membrane environments. As summarized in **Figure 2A-D**, the two plasma-membrane leaflets differ markedly in lipid-class composition, degree of unsaturation, and dominant molecular species. These quantitative features are adapted from the plasma-membrane lipidomic analysis of Lorent et al.^27^, which demonstrated enrichment of SM and PC in the outer leaflet and more highly unsaturated phospholipids in the cytoplasmic leaflet. Asymmetric lipid packing also generates transbilayer lateral pressure profiles that vary across the membrane depth. These pressure gradients can modulate conformational equilibria of membrane proteins, thereby influencing ion channels, transporters, GPCRs, and curvature-mechanosensitive proteins and receptors, independently of direct lipid binding. Lorent and colleagues further showed that this lipid-packing asymmetry is reflected in plasma-membrane protein transmembrane-domain architecture, with thinner exoplasmic regions and thicker cytoplasmic regions^27^. This relationship is schematically summarized in **Figure 2F**, linking leaflet-specific packing asymmetry with transmembrane-domain shape matching and membrane-protein organization. Reproducing both the compositional and physical asymmetry of biological membranes is therefore important for experimentally testing how membrane architecture influences membrane-protein function and protein-lipid interactions. Within this asymmetric membrane landscape, cholesterol is particularly important because it can redistribute between leaflets and modulate packing, thickness, elasticity, and curvature without changing the underlying phospholipid composition. Cholesterol therefore provides a dynamic mechanism for tuning the physical consequences of lipid asymmetry and its effects on membrane organization and protein behaviour.

### Cholesterol dynamically tunes membrane architecture

Cholesterol is a major regulator of membrane organization, continuously modulating bilayer structure and physical properties. It reduces acyl-chain motion, decreases the area per phospholipid, and increases lipid packing and bilayer thickness through intercalation between phospholipids. This condensing effect occurs across different lipid compositions, although its effect on membrane stiffness depends strongly on lipid saturation. Cholesterol markedly stiffens saturated membranes, whereas in highly unsaturated membranes it can produce only weak stiffening or even membrane softening ^30^. Thus, cholesterol regulates bilayer geometry by tuning thickness, packing, elasticity, and local curvature.

Cholesterol also helps membranes sustain the strong phospholipid imbalance between the two leaflets. Most phospholipids undergo spontaneous flip-flop only over hours to days, whereas cholesterol can move between leaflets on microsecond-to-millisecond timescales^31,32^. This rapid transbilayer motion allows cholesterol to redistribute according to differences in chemical potential and leaflet stress. Recent work on human erythrocyte membranes showed that a large phospholipid imbalance can be accommodated through an asymmetric cholesterol distribution^33^. Theoretical and simulation studies similarly show that cholesterol can move from a compressed leaflet toward a more stretched leaflet, thereby reducing differential stress generated by phospholipid number asymmetry^32^.

Cholesterol and membrane curvature are also strongly coupled. Simulations of asymmetric membranes showed that cholesterol can generate uneven curvature and that its preference for different lipid environments changes with local curvature^34^. Cholesterol contributes to spontaneous curvature indirectly through its effects on leaflet area and thickness. Models relating spontaneous curvature to leaflet area imbalance quantify this relationship as

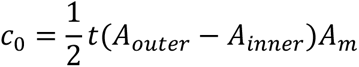

Where, t is the monolayer thickness and *A*_m_ is the average area per lipid molecule; leaflet area imbalance is a known driver of spontaneous curvature in asymmetric bilayers^35^. Cholesterol’s reduction of effective lipid area and simultaneous increase in thickness tend to bias membranes toward flatter geometries, thereby tuning the baseline curvature stress imposed by the underlying asymmetry between leaflets, consistent with cholesterol’s condensing effect on bilayer packing ^36^. Importantly, asymmetrically distributed cholesterol within the bilayer can itself generate differential stress: if cholesterol has unequal partitioning free energy between leaflets, it may accentuate or attenuate differential leaflet tension and thereby influence both spontaneous curvature and bending mechanics in ways that extend beyond composition-based curvature contributions. These effects are relevant during budding, endocytosis, exocytosis, membrane fusion, and mechano-sensation. For example, cholesterol depletion alters the activation threshold of Piezo1 and impairs curvature generation during clathrin-mediated endocytosis^30^. Thermal-gradient simulations, for example, showed that cholesterol flip-flop can become asymmetric and produce leaflet-specific enrichment^31^. Cholesterol therefore provides a fast mechanism for coupling changes in membrane composition to membrane mechanics and signaling. The primary cilium provides a physiologically relevant example of this coupling, as it is a narrow, highly curved membrane compartment that functions as a signaling center. Cholesterol organization within the ciliary membrane is essential for Hedgehog signaling through Patched1 (PTCH1) and Smoothened (SMO). Importantly, signaling depends not simply on total cholesterol but on the pool of accessible cholesterol. Hedgehog stimulation inhibits PTCH1 and increases accessible cholesterol in the ciliary membrane, whereas trapping this cholesterol suppresses downstream signaling^37^. Cholesterol can also bind directly to and activate SMO, demonstrating that sterol itself can act as a signaling molecule. Sphingomyelin helps regulate this pathway by sequestering cholesterol and limiting its accessible fraction under basal conditions ^37^. By controlling bilayer thickness, cholesterol can also modify the hydrophobic environment surrounding transmembrane domains and thereby influence hydrophobic matching. Together, these effects position cholesterol as a physical and biochemical link between lipid asymmetry, membrane curvature, membrane mechanics, protein localization, and cellular signaling^38^. These observations motivated us to test whether structurally diverse membrane proteins exhibit measurable differences in their association with curved versus flat membrane environments.

### Protein-lipid reciprocity defines structure-function cooperation

Membranes are heterogeneous and dynamic environments in which lipid organization and protein structure continuously and reciprocally influence one another. Lipid composition, packing, thickness, asymmetry and curvature shape protein conformational states and activity, whereas membrane proteins and membrane-associated proteins can reciprocally remodel local lipid organization and membrane mechanics ^1,16,39–43^. Protein-lipid reciprocity therefore provides a framework in which membrane structure and protein function emerge from their mutual coupling rather than from one-way lipid regulation.

A key manifestation of this reciprocity is hydrophobic matching, whereby the hydrophobic dimensions of proteins and bilayers are energetically coupled. Mismatch can alter transmembrane-helix orientation, membrane deformation, oligomerization and protein function, while proteins can conversely reshape local membrane thickness and packing^3,22^. Importantly, this coupling can operate at the level of individual lipid species. Lipid-specific interactomics has revealed selective associations of phosphatidylethanolamine with membrane-organizing proteins such as MIC60 and NDC1, linking lipid identity to protein-complex organization in curved membranes^44^. Similarly, perturbation of ER lipid composition can alter membrane-dependent signalling, as exemplified by changes in phosphatidylethanolamine abundance following DGAT2 inhibition and consequent suppression of SREBP1 cleavage ^45,46^. These observations support a dynamic feedback model in which lipid composition influences protein activity, while proteins reciprocally modify the physical and functional state of the surrounding membrane. This reciprocity becomes particularly apparent when disease-associated lipid remodelling changes the physical environment of membrane proteins. Lipidomic analysis of mammary tumours, for example, revealed marked enrichment of phosphatidylserine and ether-linked/plasmalogen PS species, including approximately 19-fold enrichment of PS(O-18:0/18:0) and greater than 12-fold enrichment of several plasmalogen PS species. Many of these species contain highly unsaturated fatty acyl chains, which can influence membrane packing, flexibility and lateral organization. Although increased PS abundance promotes MERTK-dependent signalling and tumour-associated macrophage activity, such remodelling may additionally alter membrane charge, packing and curvature, thereby modifying the physical landscape experienced by membrane proteins. Thus, disease-associated lipidomic changes can propagate from lipid composition to membrane biophysics and, ultimately, to protein organization and signalling.

Protein-lipid reciprocity is not restricted to integral membrane proteins. Soluble proteins can transiently acquire membrane-dependent conformations and activities, while their membrane association can subsequently perturb lipid organization. Cholesterol-dependent cytolysins provide a classical example in which recognition of accessible cholesterol promotes protein conformational changes and pore formation, with the resulting oligomers disrupting membrane organization. Our previously published studies of the *Mycobacterium tuberculosis* secretory protein MPT63 provide a related example (**Figure 3A**). MPT63 undergoes a pH-dependent β-sheet-to-α-helical conformational transition involving a chameleon sequence, converting a membrane-inactive protein into a pore-forming oligomer capable of inducing macrophage cell death ^47–50^. Here, environmental conditions determine the protein conformational state, which subsequently becomes coupled to membrane disruption^51^.

**Figure 3.**
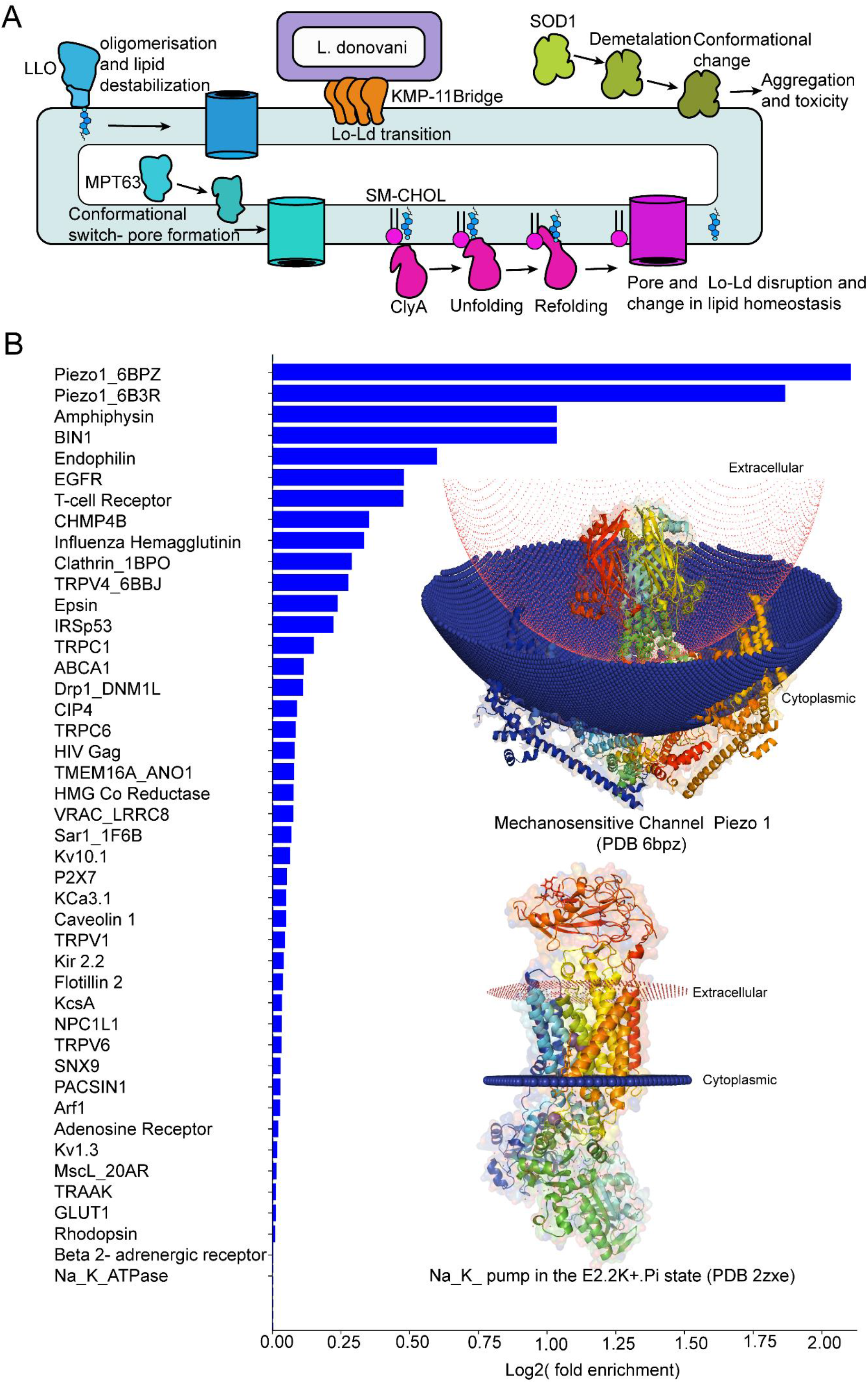
Representative examples of lipid-protein reciprocal coupling and curvature-dependent membrane association. **(A)** Analytical cell model mapping diverse host-pathogen interactions that disrupt membrane homeostasis. Toxic pathways include listeriolysin O (LLO) oligomerization and lipid destabilization, *Leishmania donovani* binding via a KMP-11 bridge to trigger a liquid-ordered to liquid-disordered (Lo-Ld) transition, bacterial MPT63 pore formation, SOD1 demetalation inducing toxic aggregations, and ClyA unfolding/refolding sequences that form transmembrane pores across SM-CHOL rich targets. **(B)** Relative fold enrichment chart measuring specialized protein distributions across highly curved geometries. The mechanosensitive channel Piezo1 exhibits the highest affinity for highly curved membranes, followed by curvature-generating elements (amphiphysin, BIN1, endophilin, EGFR). Insets contrast a 3D structural model of the curved Piezo1 mechanosensitive channel (PDB: 6BPZ) sitting inside a matching cytoplasmic indentation against a flat membrane baseline typified by the Na-K pump in the E2.2K+.Pi state (PDB: 2ZXE).

KMP-11 from *Leishmania donovani* illustrates an even more direct form of reciprocal coupling in which protein binding alters lipid molecular conformation and the resulting membrane state feeds back on protein behaviour. KMP-11 oligomers bridge parasite and host membranes in a sterol-dependent manner, facilitating cholesterol transfer and subsequent host-membrane phase transitions during parasite invasion^52–55^. At the molecular level, KMP-11 binding to phospholipid membranes increases the population of gauche lipid-chain conformers at the expense of the ordered trans conformers, as detected by FTIR analysis. This trans-to-gauche transition increases acyl-chain disorder and promotes the transition towards a more fluid liquid-disordered membrane state^52^. Thus, KMP-11 does not simply sense a pre-existing membrane phase; its interaction actively changes lipid-chain conformational order. Conversely, the resulting membrane state influences KMP-11 binding: KMP-11 displays stronger association with ordered/gel-like membrane states but weaker retention following the protein-induced transition towards the liquid-disordered state, facilitating parasite detachment after cholesterol transfer^52^. Our previously reported KMP-11 studies therefore provide a molecular example of a feedback loop in which protein binding drives a lipid conformational transition, while the altered lipid state subsequently regulates protein-membrane affinity and function. The same principle may operate in pathological transitions of soluble proteins. Superoxide dismutase 1 (SOD1), for example, can acquire membrane-dependent behaviour when its metal-cofactor state is perturbed. Zinc deficiency alters SOD1 conformation and membrane association through changes involving flexible loop regions, promoting lipid-induced aggregation in an in vitro model of amyotrophic lateral sclerosis^56^. In this setting, protein cofactors, conformational stability and membrane interactions become interconnected variables, suggesting that membrane environments can participate in the transition between functional and pathological protein states(**Figure 3A**).

Our previously reported Cytolysin A (ClyA) study provides a particularly direct demonstration of this concept, in which the membrane actively promotes protein conformational maturation. Rather than functioning simply as a surface for toxin binding, a sphingomyelin-cholesterol environment acts as a lipid chaperone that facilitates the conformational transitions required for pore assembly. Sphingomyelin-cholesterol synergy promotes membrane-associated unfolding of the β-tongue region into a reactive molten-globule-like intermediate and subsequently favours refolding into the α-helical architecture required for productive oligomerization and pore formation ^57^. Once assembled, ClyA pores can also alter the surrounding lipid environment by redistributing sphingomyelin and cholesterol, changing liquid-ordered and liquid-disordered membrane organization, and disrupting membrane homeostasis^57^. Thus, lipid organization controls protein conformational maturation, whereas protein assembly feeds back to reorganize the lipid landscape. Together, these examples support protein-lipid reciprocity as a multiscale principle of membrane organization (**Figure 3A**) and provide a mechanistic context for the curvature-dependent protein association analyzed below.

### Membrane Curvature Preferentially Tunes Protein-Membrane Association

Curvature preference therefore cannot be understood from membrane geometry alone, because changes in curvature are accompanied by changes in lipid packing and in the hydrophobic environment experienced by membrane proteins. This coupling brings hydrophobic matching into focus as an additional physical principle through which membrane architecture can influence protein organization and function. Membrane proteins interact with lipid bilayers within a dynamic physical environment in which membrane composition, packing, geometry, and curvature can influence protein localization and function. Although many membrane-associated proteins bind to lipid bilayers, their membrane association is not necessarily independent of membrane geometry. To test this relationship quantitatively, we analyzed 44 membrane proteins using membrane orientation information from the OPM (Orientation of Proteins in Membranes) framework ^84,85^ and quantified their relative association with curved and flat membrane environments. The resulting fold enrichment represents the ratio of curved-membrane to flat-membrane association and therefore provides a measure of curvature preference, rather than absolute membrane-binding affinity.

Our analysis revealed substantial heterogeneity in curvature-dependent membrane association across the 44 proteins (**Figure 3B**, **Table 1**). A value greater than one indicates preferential association with the curved membrane, whereas a value close to one indicates little difference between curved and flat membranes. Several proteins with established roles in membrane curvature sensing or generation occupied the upper end of the ranking, whereas many other membrane proteins showed weaker curvature preference. Thus, curvature sensitivity appears to vary substantially among membrane proteins rather than representing a universal property of membrane association.

**Table 1.**
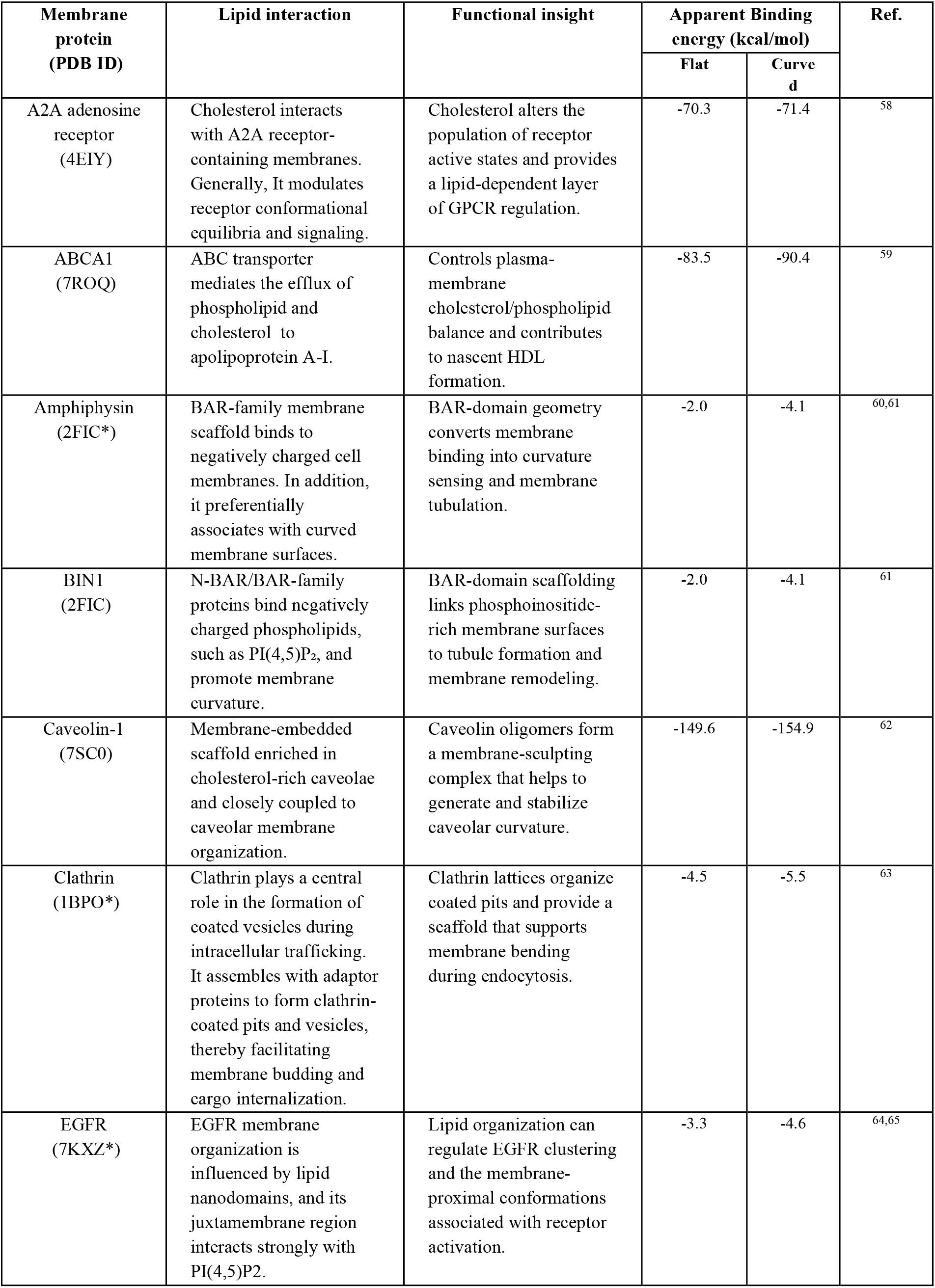
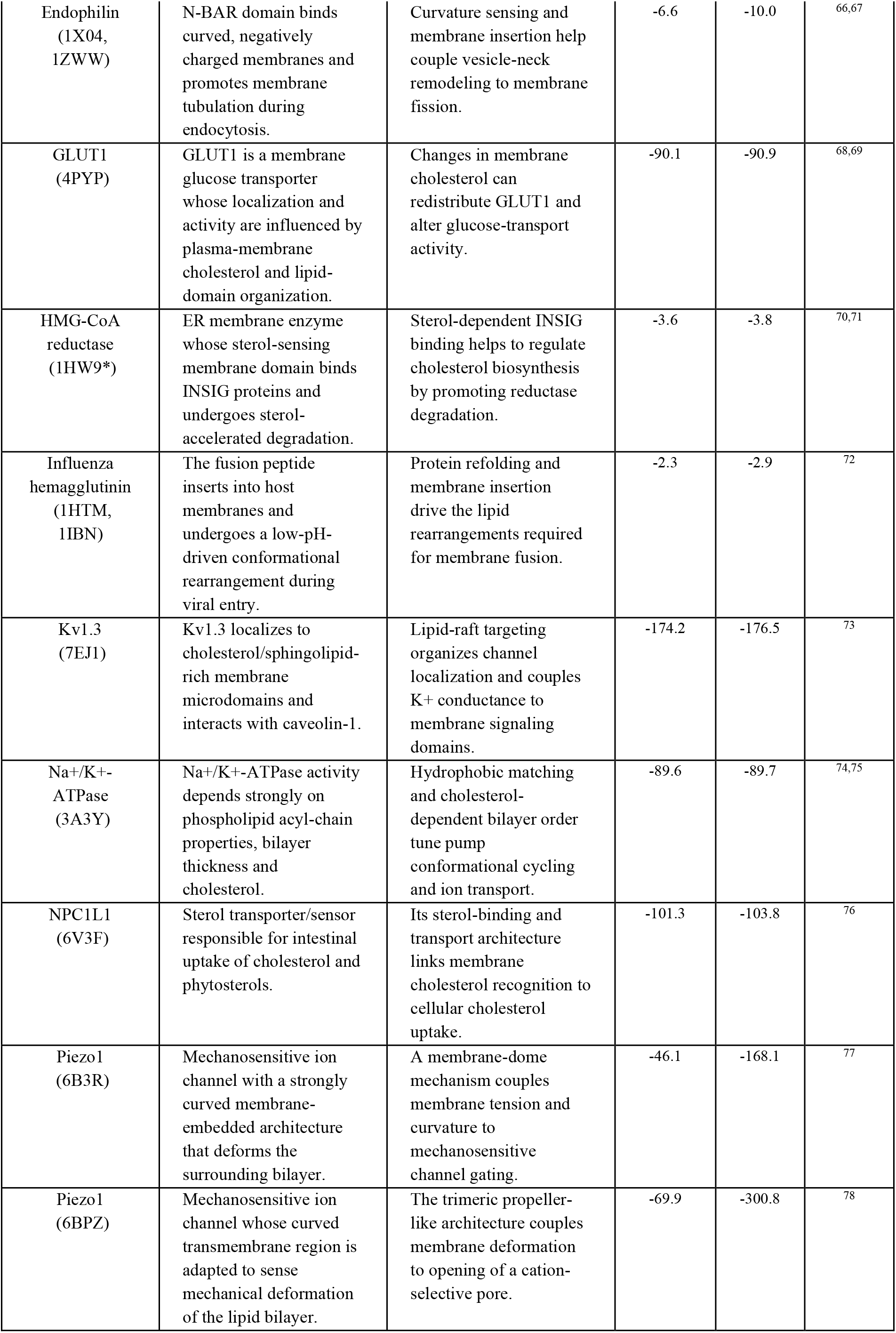
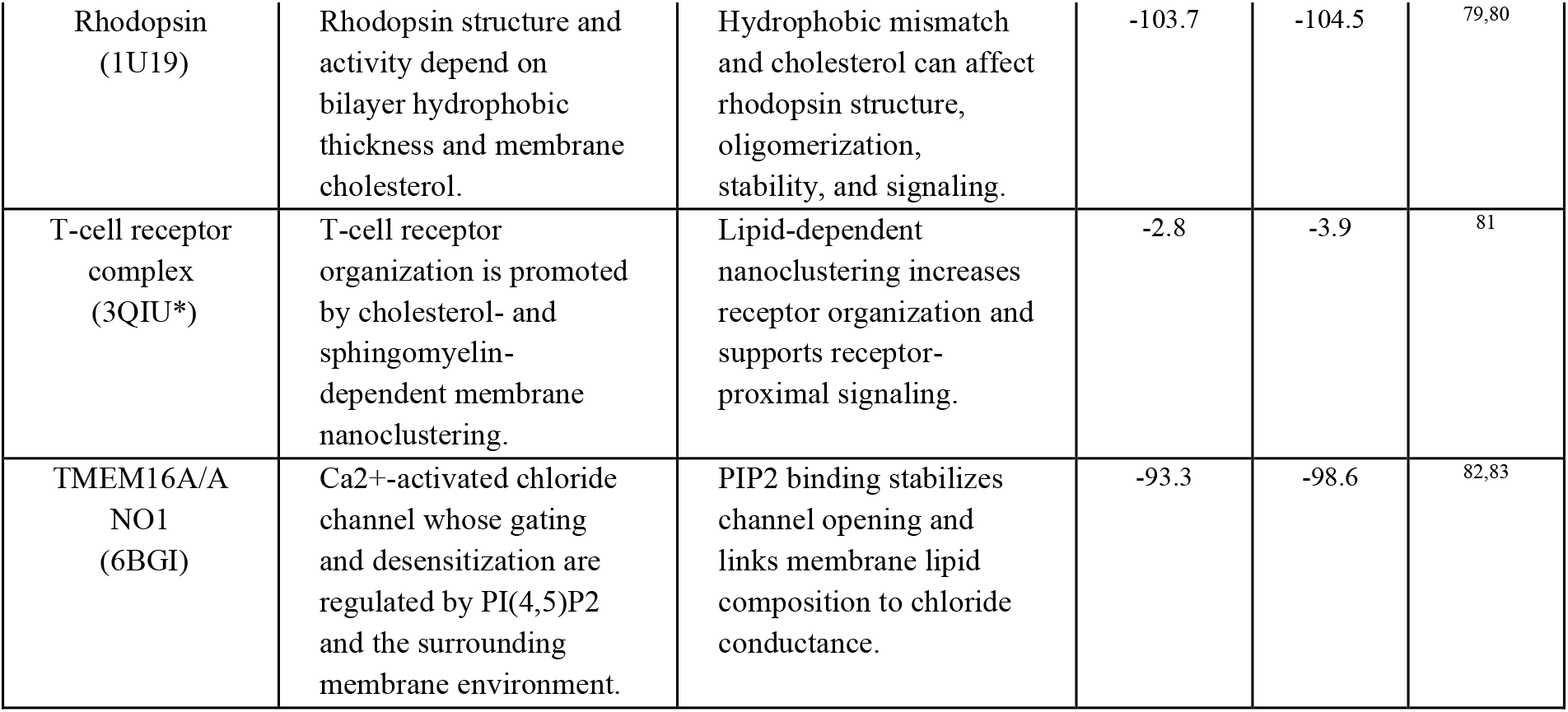
A quantitative analysis of diverse membrane protein binding with flat and planar lipid bilayer, their respective lipid interactions, and their functional insights.

The strongest curvature preference was observed for the two Piezo1 structures, followed by amphiphysin and BIN1, with endophilin also showing substantial enrichment. Piezo1 is particularly notable because its trimeric architecture forms a pronounced curved structure that deforms the surrounding bilayer and undergoes force-dependent conformational changes^86,87^. Consistent with this structural relationship, membrane curvature regulates the distribution of Piezo1 in living cells, with Piezo1 enriched in nanoscale membrane invaginations and depleted from highly curved membrane protrusions^86^. The strong curved-to-flat enrichment observed for Piezo1 in our analysis is therefore consistent with the established coupling between Piezo1 architecture and membrane geometry.

This strong curvature preference may also have disease relevance. Piezo1 couples membrane mechanics to ion-channel activity through force-dependent opening of its central pore, allowing cation influx, particularly Ca²⁺. Its activity is influenced by membrane lipids such as cholesterol and PIP₂, which contribute to its mechanosensitivity and conformational regulation^87^. Dysregulated Piezo1-mediated mechanotransduction has been associated with cancer-cell migration and invasion, suggesting that altered membrane mechanics and curvature sensing may contribute to tumor progression^88^.

A similar relationship is evident for BAR-domain proteins. Amphiphysin contains a crescent-shaped BAR domain that recognizes membrane curvature and can promote membrane deformation ^60^. BIN1 also possesses a curved BAR domain with a positively charged membrane-interacting surface that contributes to membrane binding and tubulation^61^. Endophilin provides another example, as its BAR domain and amphipathic helices cooperate to sense and generate membrane curvature during membrane remodeling ^66,67^. The high curved-membrane enrichment identified for these proteins in our analysis therefore agrees with their established roles in curvature sensing and membrane remodeling.

The disease relevance of amphiphysin further emphasizes the importance of curvature-dependent membrane organization. Amphiphysin is involved in neuronal membrane remodeling and endocytosis, and autoantibodies against amphiphysin are strongly associated with stiff-person syndrome and related neurological disorders. In addition, dysregulation of BAR-domain proteins can affect membrane and cytoskeletal remodeling processes involved in cancer-cell migration and invasion^89,90^. Thus, proteins with strong curvature preference may have pathological consequences when their membrane localization or remodeling activity is disrupted.

Importantly, not all proteins showing curvature preference are canonical curvature sensors. EGFR, for example, interacts with PIP₂ through its intracellular juxtamembrane region, where these interactions contribute to receptor organization and activation^64^. Similarly, T-cell receptor signaling is influenced by membrane topology and mechanical forces, with local membrane bending proposed to contribute to receptor triggering^91^. These examples indicate that curvature preference can arise from protein architecture, lipid interactions, or membrane mechanics even when curvature sensing is not the primary function of the protein.

EGFR also illustrates the connection between membrane organization and cancer. Although EGFR is not a classical curvature-sensing protein, its interactions with PIP₂ influence receptor organization, clustering, and activation, while receptor stimulation promotes membrane invagination and endocytic trafficking^64^. Because abnormal EGFR signaling promotes cell proliferation, survival, and migration, dysregulation of this pathway is strongly associated with cancer progression^92^. Thus, membrane geometry and lipid organization may influence disease-associated signaling even in proteins that do not directly function as curvature sensors.

CIP4 and Caveolin-1 further illustrate the connection between membrane association and curvature generation. CIP4 is an F-BAR protein that can bind membranes and organize into assemblies capable of generating membrane tubules^93^. Caveolin-1 interacts with cholesterol-rich membranes and can promote membrane curvature, with the extent of curvature influenced by membrane cholesterol ^94^. CIP4 also connects membrane remodeling with the actin cytoskeleton and has been implicated in cancer-cell migration, invadopodia formation, and invasion^95^. Together, these examples emphasize that curvature preference can emerge from the combined effects of lipid recognition, protein geometry, membrane insertion, oligomerization, and cytoskeletal interactions, with potential consequences for disease-associated membrane remodeling.

Our ranking further indicates that membrane association and curvature preference are related but distinct properties. Proteins such as Piezo1, amphiphysin, BIN1, and endophilin show strong curvature preference consistent with their established roles in membrane deformation or curvature recognition. In contrast, proteins with lower curved-to-flat enrichment may still associate efficiently with membranes but show little discrimination between the two membrane geometries. Importantly, a low enrichment value should not be interpreted as weak membrane binding; rather, it indicates a weaker preference for the curved membrane relative to the flat reference.

The upper portion of the quantitative ranking further highlights these differences. The two Piezo1 structures show log₂ fold-enrichment values of approximately 2.1 and 1.9, corresponding to roughly 4.3- and 3.7-fold greater association with the curved membrane, respectively. Amphiphysin and BIN1 show approximately two-fold enrichment, whereas endophilin shows approximately 1.5-fold enrichment. These differences indicate that curvature preference exists on a continuum rather than as a simple binary property.

Overall, the analysis of 44 membrane proteins demonstrates that curvature preference is heterogeneous across membrane proteins and is particularly pronounced among several established curvature-sensing or curvature-generating proteins. Rather than being determined solely by membrane-binding capacity, curvature recognition likely reflects the combined effects of protein architecture, lipid interactions, membrane insertion, hydrophobic mismatch, and the physical geometry of the protein-membrane interface. Importantly, these physical interactions may have disease relevance because altered membrane curvature and remodeling can affect mechanosensation, receptor signaling, membrane trafficking, cytoskeletal organization, and cancer-cell invasion. Our curved-to-flat enrichment analysis therefore provides a quantitative framework for distinguishing general membrane association from curvature-selective membrane association and for linking membrane geometry with protein function and disease-associated membrane remodeling.

### Hydrophobic landscapes: connecting membrane topology with membrane protein design

Hydrophobic matching is an important physical principle that connects membrane structure with membrane-protein organization and function. Hydrophobic mismatch arises when the hydrophobic length of a transmembrane domain differs from the hydrophobic thickness of the surrounding lipid bilayer. Because lipid asymmetry, acyl-chain composition, cholesterol content, and membrane curvature can alter local bilayer thickness and packing, changes in membrane architecture can directly change the extent of mismatch experienced by a membrane protein ^3^. To reduce the associated energetic cost, the membrane-protein system can respond through local membrane thinning or thickening, transmembrane-helix tilting, protein clustering or oligomerization, and conformational rearrangement of the protein (**Figure 4A**)^3,95^. In cells, biological membranes differ in hydrophobic thickness owing to organelle-specific lipid composition. The endoplasmic reticulum (ER), which is lower in cholesterol and sphingolipids and enriched in unsaturated phospholipids, forms one of the thinnest cellular membranes, whereas the Golgi exhibits intermediate thickness and the cholesterol- and sphingolipid-rich plasma membrane is the thickest membrane in the secretory pathway. Membrane proteins have co-evolved with these distinct lipid environments, such that the hydrophobic length of their transmembrane domains closely matches the thickness of their resident membrane. Accordingly, ER-resident proteins typically possess shorter transmembrane helices, whereas Golgi and plasma membrane proteins contain progressively longer hydrophobic segments, thereby minimizing hydrophobic mismatch and contributing to organelle-specific protein sorting and retention ^25,96,97^. Perturbations in membrane lipid composition or mutations that alter transmembrane hydrophobic length can disrupt this hydrophobic matching, resulting in protein mislocalization, altered conformational stability, and impaired function.

**Figure 4.**
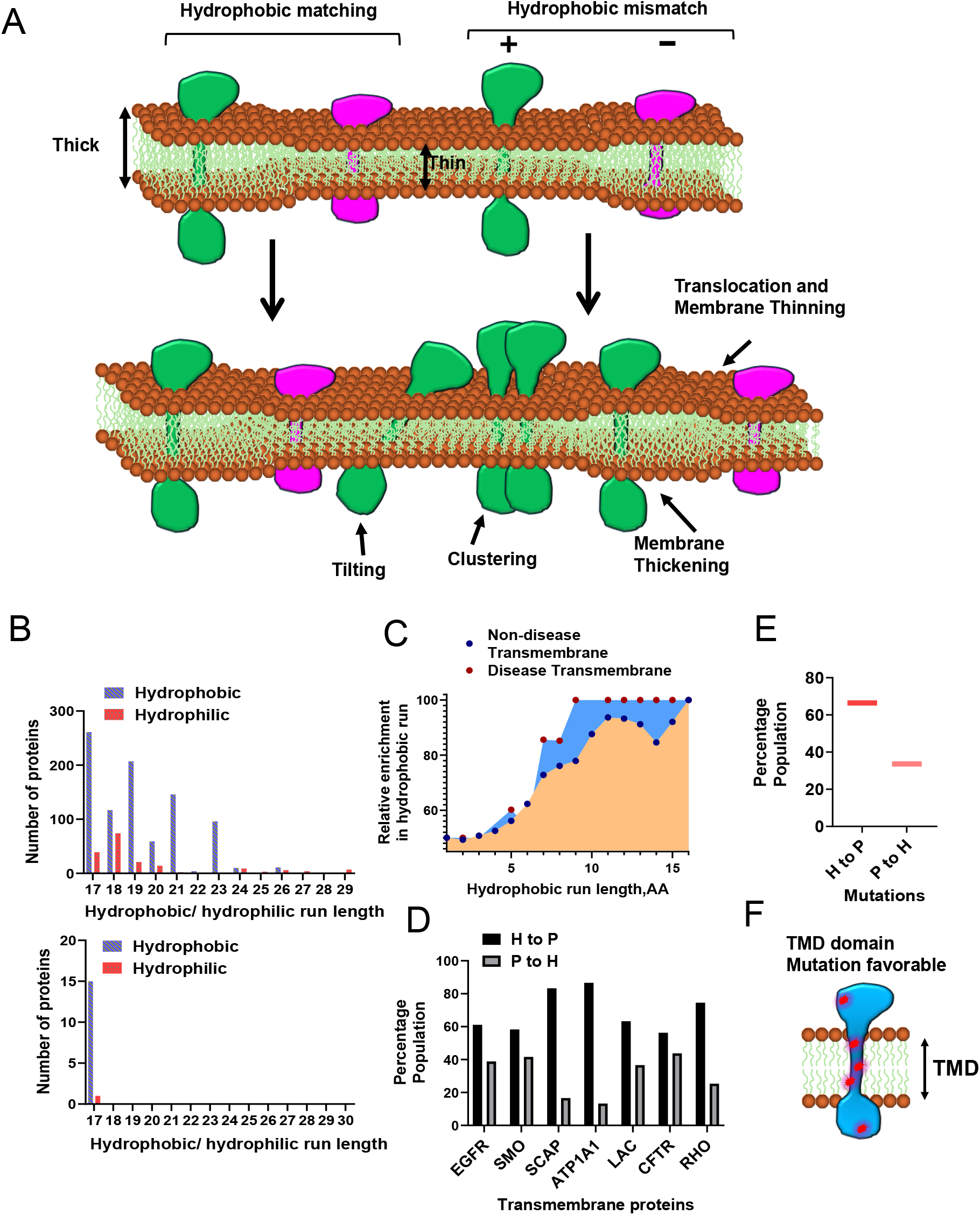
Hydrophobic mismatch, hydrophobic-run organization, and mutation patterns in transmembrane domains (TMDs). **(A)** Schematic illustration of hydrophobic matching and mismatch between an α-helical transmembrane protein and lipid bilayers of different thicknesses. Optimal matching minimizes free-energy cost, whereas positive (+) and negative (−) mismatch can induce local adaptations such as protein clustering, membrane thickening or thinning, helix tilting, or protein translocation. **(B)** Distribution of consecutive hydrophobic (blue) and hydrophilic (red) amino-acid run lengths within protein sequences, showing the organization of sequence runs across large protein datasets. **(C)** Comparison of hydrophobic-run length distributions between non-disease-associated transmembrane proteins (blue) and disease-associated transmembrane proteins (red), showing enrichment of longer hydrophobic runs in the neutral dataset and shorter runs in the disease-associated dataset. **(D)** Frequencies of hydrophobic-to-polar (H→P) and polar-to-hydrophobic (P→H) substitutions in representative disease-associated membrane proteins (EGFR, SMO, SCAP, ATP1A1, LAC, CFTR, and RHO). **(E)** Aggregate distribution of mutation types across the analyzed transmembrane proteins, showing a higher prevalence of H→P substitutions. **(F)** Schematic representation of a TMD containing a disruptive mutation, such as an H→P substitution (red stars), which introduces polar residues into the hydrophobic core and perturbs native helix packing.

Rhodopsin provides a well-studied example of this effect. Rhodopsin is a seven-transmembrane G-protein-coupled receptor in retinal photoreceptor cells that initiates visual signaling following light activation. Its conformational equilibrium is sensitive to the hydrophobic thickness of the surrounding membrane. Changes in bilayer thickness alter the helical organization of rhodopsin and shift the equilibrium between the metarhodopsin I and signaling-active metarhodopsin II states^98^. Hydrophobic mismatch can also promote rhodopsin oligomerization, whereas a bilayer hydrophobic thickness of approximately 27 ± 1 Å. produces minimal membrane perturbation ^79^. Membrane curvature and hydrophobic forces further influence rhodopsin oligomerization and activity^99^. These studies illustrate a broader principle in which hydrophobic mismatch can regulate the orientation, stability, clustering, conformation, localization, and activity of transmembrane proteins.

At the sequence level, hydrophobic matching is closely related to the organization of hydrophobic residues within membrane proteins. Transmembrane proteins usually contain continuous stretches of hydrophobic amino acids, known as hydrophobic runs, that help their membrane-spanning regions interact with the hydrophobic core of the lipid bilayer. Previous statistical analyses of proteins of known structure showed that long hydrophobic runs are strongly suppressed in soluble globular proteins but are considerably more common in membrane and cell-surface proteins^100,101^. In one analysis, soluble proteins did not contain hydrophobic runs longer than 16 residues, whereas longer hydrophobic runs were found in membrane-associated proteins and were frequently associated with transmembrane segments (**Figure 4B**).

Importantly, long hydrophobic runs are not restricted to proteins that permanently reside within membranes. Soluble proteins that subsequently associate with or insert into membranes can also contain unusually long hydrophobic motifs. Cytolysin A (ClyA), a soluble pore-forming toxin, contains a 16-residue hydrophobic run within its β-tongue region, which undergoes a major structural rearrangement during membrane insertion^57^. Colicin 1A, another pore-forming protein that is soluble before membrane association, also contains an extended hydrophobic segment within its membrane-interacting region^57^. In ClyA, sphingomyelin and cholesterol were proposed to create a favorable lipid environment for the transition from the soluble state to the membrane-inserted pore-forming state. These examples suggest that long hydrophobic runs can serve as functional membrane-interacting motifs even in proteins that are not constitutively embedded in the bilayer.

The length and continuity of these hydrophobic runs may therefore be important for maintaining an appropriate match between the protein and its surrounding membrane. To test this relationship at the sequence level, we performed a bioinformatic analysis using structural protein datasets and disease-associated variants. Our analysis comparing SCOP-derived neutral proteins with proteins containing disease-associated mutations from the mutHTP database showed that longer hydrophobic runs occur more frequently in the neutral dataset, whereas disease-associated proteins are enriched in shorter hydrophobic runs (**Figure 4C**). In our ClyA-associated analysis, disease-associated variants with very long hydrophobic runs were strongly reduced or absent^57^. These findings suggest that missense mutations can disrupt continuous hydrophobic sequences and reduce their effective run length.

The chemical nature of these substitutions provides a possible physical explanation for this trend. A hydrophobic-to-polar substitution within a membrane-spanning hydrophobic run can shorten the effective hydrophobic region of the protein and increase the energetic penalty of placing that segment within the bilayer. In contrast, a polar-to-hydrophobic substitution can extend or alter the hydrophobic region and thereby change its interaction with the surrounding membrane. Our mutation analysis summarized in **Figure 4D-F** indicates that, for several representative membrane proteins, disease-associated substitutions occur frequently within hydrophobic-run regions, with hydrophobic-to-polar substitutions occurring more often than the reverse polar-to-hydrophobic substitutions^57^. Such mutations can change membrane insertion, helix orientation, oligomerization, conformational stability, or protein localization.

Together, our statistical analyses suggest an important connection between hydrophobic-run disruption and hydrophobic mismatch. A missense mutation that converts a hydrophobic residue to a polar residue can interrupt a continuous hydrophobic sequence and reduce the effective hydrophobic length of a transmembrane segment. The resulting protein may no longer match the hydrophobic thickness of its native membrane as efficiently, increasing the energetic cost of membrane insertion and potentially altering protein structure or function. The effect of such a mutation will also depend on the physical properties of the surrounding membrane, because lipid composition, leaflet asymmetry, cholesterol content, membrane curvature, and membrane mechanics all influence local bilayer thickness and packing. Thus, hydrophobic mismatch can arise either from changes in the membrane or from changes in the hydrophobic organization of the protein itself.

This connection is particularly relevant because membrane architecture is dynamic rather than fixed. Changes in lipid asymmetry or curvature can alter the local hydrophobic environment even when the protein sequence remains unchanged, whereas disease-associated mutations can modify hydrophobic-run length even when the membrane composition remains constant. Together, these effects establish a close relationship between membrane lipid architecture, hydrophobic-run organization, protein topology, and disease-associated dysfunction.

The functional importance of this coupling is also evident in proteins whose activities depend on specific lipid interactions. Our previous ClyA study provides one example in which sphingomyelin and cholesterol promote the transition from a soluble state to a membrane-inserted pore-forming state^57^. Similar lipid-dependent mechanisms have been described in numerous membrane proteins, including EGFR, CFTR, ATP1A1, Smoothened, and rhodopsin, where alterations in membrane composition remodel conformational equilibria and signaling outputs. Smoothened is particularly relevant because cholesterol regulates Hedgehog signaling through interactions with its cysteine-rich and transmembrane regions, whereas sphingomyelin can modulate cholesterol accessibility. These examples further illustrate how hydrophobic sequence organization and the surrounding lipid environment can work together to regulate membrane-protein structure, signaling, and disease-associated dysfunction. Resolving these context-dependent relationships requires experimental approaches that can independently manipulate membrane composition and physical properties while directly measuring protein behavior. Biomimetic membrane systems provide such controlled environments, allowing lipid asymmetry, curvature, thickness and mechanics to be systematically varied to experimentally test their contributions to membrane-protein coupling.

### Decoding membrane architecture: sensors and biomimetic systems

Model membrane systems provide controlled platforms for testing how lipid organization, membrane mechanics and geometry influence protein-lipid interactions (**Figure 5**). Liposomes are among the most widely used models: SUVs and LUVs are useful for studying membrane binding, transport, fusion and curvature, whereas GUVs enable direct visualization of phase separation and membrane mechanics (**Figure 5A**). GPMVs retain much of the lipid and protein complexity of native plasma membranes and are useful for studying membrane domains and protein partitioning^102^. However, vesicles can exhibit heterogeneity in size and composition, while GPMVs lack active cytoskeletal organization. Micropipette aspiration provides quantitative measurements of GUV tension and elasticity but is relatively low-throughput and requires individual vesicle manipulation.

**Figure 5.**
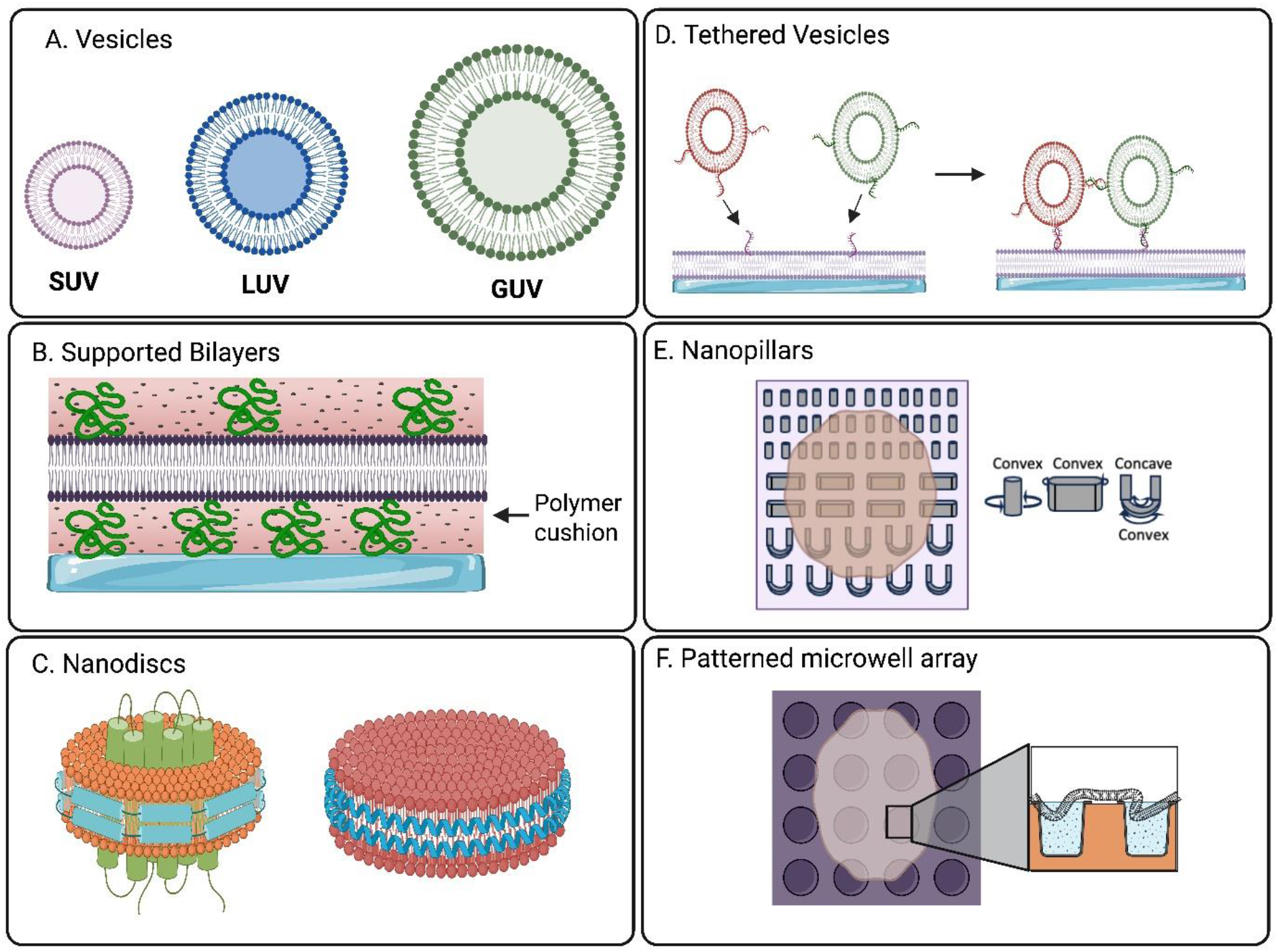
Representative biomimetic membrane platforms for studying membrane organization and lipid-protein interactions. **(A)** Lipid vesicles of different sizes, including small unilamellar vesicles (SUVs), large unilamellar vesicles (LUVs), and giant unilamellar vesicles (GUVs), provide tunable membrane systems for studying lipid composition, curvature, and protein association. **(B)** Supported lipid bilayers, including polymer-cushioned configurations, provide planar membrane surfaces while reducing direct substrate-membrane interactions. **(C)** Nanodiscs enable membrane proteins and defined lipid compositions to be reconstituted within nanoscale bilayer patches stabilized by scaffold proteins or related supports. **(D)** Tethered vesicle systems immobilize vesicles at supported membrane surfaces, allowing controlled studies of vesicle-vesicle and vesicle-membrane interactions. **(E)** Nanopillar platforms impose defined local membrane geometries, including convex and concave curvature, for investigating curvature-dependent protein localization and membrane remodeling. **(F)** Patterned microwell arrays support suspended or pore-spanning membrane configurations, providing spatially defined membrane regions for optical, mechanical, and functional measurements.

Supported lipid bilayers (SLBs) provide stable planar membranes compatible with fluorescence microscopy, AFM and studies of lateral diffusion and protein-lipid interactions (**Figure 5B**). Their major limitation is interaction with the solid substrate, which can restrict lipid and transmembrane-protein mobility; polymer cushions can reduce but not eliminate these effects ^103^. Nanodiscs provide small, soluble bilayer patches that are particularly useful for structural and biochemical studies of membrane proteins (**Figure 5C**), although their finite size and surrounding scaffold can influence membrane properties and protein conformations ^104^.

Tethered-vesicle systems enable controlled measurements of vesicle docking and membrane fusion, particularly for SNARE-mediated fusion (**Figure 5D**), but surface attachment and simplified reconstitution can alter vesicle mobility and fusion kinetics ^105^. Nanopillar systems impose defined nanoscale curvature on cellular membranes and enable studies of curvature-dependent protein recruitment (**Figure 5E**); however, substrate adhesion, membrane tension and cytoskeletal responses can complicate interpretation of curvature-specific effects^106^. Patterned microwell arrays provide another approach for generating suspended membranes with controlled geometry while minimizing direct substrate contact (**Figure 5F**).

Collectively, these systems illustrate the trade-off between experimental control and biological complexity in studies of lipid-protein coupling. No single platform fully reproduces the compositional, mechanical and dynamic properties of cellular membranes, emphasizing the value of complementary model systems for testing specific aspects of lipid-protein coupling. Complementary tools are therefore required not only to reconstruct membrane environments but also to determine where specific lipid species are located and how their distributions change in space and time. Molecular lipid sensors provide this additional level of information by enabling leaflet-specific and spatially resolved measurements of membrane composition and lipid accessibility.

### Molecular Sensors for Asymmetric Lipids and Distinct Cholesterol Pools

The asymmetric and dynamic organization of membrane lipids makes direct visualization essential for determining how local lipid organization influences signalling and lipid-protein interactions. Although bulk lipid measurements provide information on overall membrane composition, they cannot readily resolve leaflet-specific localization, local lipid organization or rapid lipid redistribution. Protein-based lipid sensors are therefore valuable tools because naturally occurring toxins and lipid-binding proteins can recognize specific membrane lipids with high selectivity, whereas engineered variants can provide quantitative and spatially resolved measurements in living cells. For example, lysenin recognizes sphingomyelin (SM), annexin V detects phosphatidylserine (PS), and duramycin binds phosphatidylethanolamine (PE). Lipid-binding domains provide additional molecular specificity, including the PLCδ1-PH domain for PI(4,5)P₂, the TAPP1-PH domain for PI(3,4)P₂, the GRP1-PH domain for PI(3,4,5)P₃, Lact-C2 for PS and the Spo20 phosphatidic-acid-binding domain for phosphatidic acid (**Figure 6A**). These probes enable spatial redistribution of specific lipids to be directly related to signalling events and membrane-protein recruitment.

**Figure 6.**
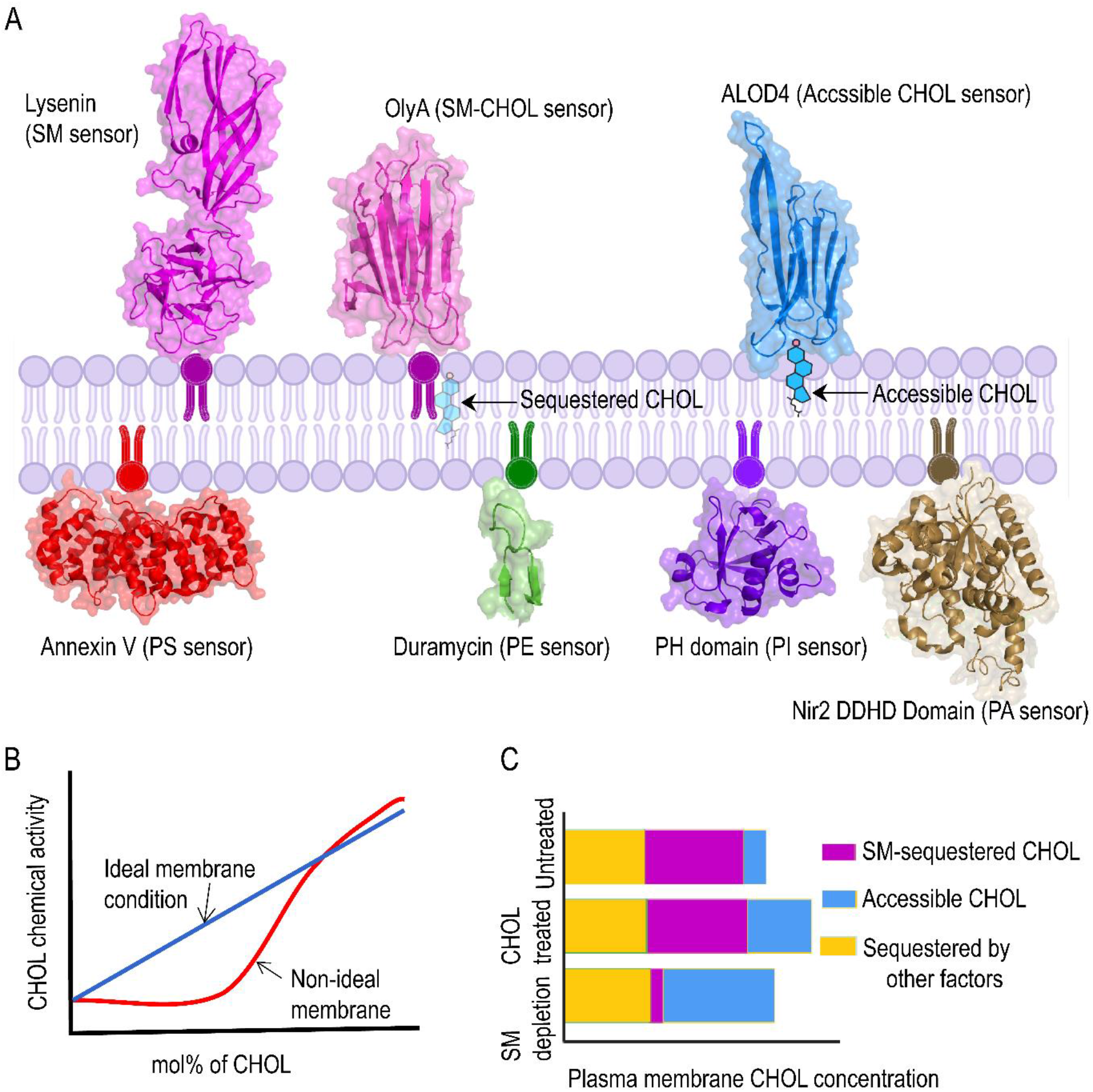
Molecular sensors for asymmetric membrane lipids and distinct cholesterol pools. **(A)** Representative lipid-sensing proteins and probes used to detect specific membrane lipids and cholesterol states, including lysenin for sphingomyelin (SM), OlyA for SM-cholesterol assemblies, ALOD4 for accessible cholesterol, Annexin V for phosphatidylserine (PS), duramycin for phosphatidylethanolamine (PE), PH domains for phosphoinositides (PI), and the Nir2 DDHD domain for phosphatidic acid (PA). The schematic also distinguishes accessible cholesterol from cholesterol sequestered within SM-rich membrane environments. **(B)** Conceptual relationship between membrane cholesterol concentration and cholesterol chemical activity, illustrating ideal and non-ideal membrane behaviour. **(C)** Schematic representation of plasma-membrane cholesterol partitioning into SM-sequestered cholesterol, accessible cholesterol, and cholesterol sequestered by other factors, together with redistribution of these pools following cholesterol treatment or sphingomyelin depletion.

Cholesterol requires particular consideration because it exists in functionally distinct pools that differ in accessibility to sensing proteins and in their availability for cellular transport and metabolism. Das and colleagues distinguished an accessible cholesterol pool, an SM-sequestered pool that becomes accessible following sphingomyelin hydrolysis, and a residual pool that remains inaccessible even after SM depletion (**Figure 6 B-C**)^107^. Importantly, cholesterol sensing at the plasma membrane and endoplasmic reticulum (ER) is highly nonlinear, indicating that cholesterol availability is governed by cooperative interactions and threshold behaviour rather than by a simple linear relationship with total membrane cholesterol. A critical cholesterol threshold has been identified at both the plasma membrane and ER, above which cholesterol-dependent signalling and transport processes, including toxin binding at the plasma membrane and SCAP-SREBP-dependent cholesterol homeostasis at the ER, are activated^107,108^. Toxin-based sensors have therefore provided a powerful means of distinguishing cholesterol pools on the basis of their accessibility rather than simply their abundance. The cholesterol-binding D4 domain of perfringolysin O (PFO) has been engineered into probes such as D4H, as well as affinity-tuned DAN-D4 and NR3-D4 variants, which detect cholesterol over different concentration ranges^109^. Complementary extracellular D4 probes and cytosolic D4H sensors can further distinguish cholesterol in the exoplasmic and cytoplasmic leaflets of the plasma membrane, respectively, while interactions with PS contribute to cholesterol retention in the cytoplasmic leaflet^110^. Clickable cholesterol analogues and fluorescent sterols such as BODIPY- and TopFluor-cholesterol provide complementary approaches for monitoring sterol trafficking and redistribution in living cells^111–115^. More recently, lipid sensing has expanded beyond individual lipid species to the detection of defined lipid-lipid assemblies. Ostreolysin A (OlyA), derived from a fungal pore-forming protein, selectively recognizes SM-cholesterol complexes, and structural studies have directly visualized this interaction within lipid bilayers ^116,117^. Such probes enable cholesterol organized within SM-rich assemblies to be distinguished from cholesterol defined solely by its chemical abundance.

Since membrane lipid organization is highly dynamic, current approaches are increasingly moving towards real-time, quantitative and single-molecule measurements. A genetically encoded FRET-based sensor has recently been developed to monitor SM-cholesterol organization and plasma-membrane mechanical properties in living cells under changing mechanical and environmental conditions^118^. Together with single-molecule and super-resolution imaging approaches^119^, these tools enable lipid redistribution and nanoscale organization to be related to membrane mechanics and cellular signalling. Nevertheless, selective probes remain unavailable for several abundant membrane lipids. In particular, robust sensors for phosphatidylcholine (PC), one of the most abundant phospholipids in animal membranes, would substantially expand our ability to monitor leaflet organization, lipid redistribution and lipid-protein interactions. Future probe development should therefore move beyond the recognition of individual lipid species towards quantitative sensing of lipid-lipid assemblies and lipid-protein interactions. Such approaches could provide a more integrated view of membrane asymmetry, lipid organization and the dynamic regulation of cellular membrane function. These molecular sensors provide powerful tools for resolving lipid organization, but directly testing lipid-protein reciprocal coupling requires membrane platforms that combine controlled asymmetry and geometry with optical and functional measurements. Suspended pore-spanning membranes offer a promising route toward this integration.

### Suspended Pore-Spanning Membranes for High-Throughput Studies of Membrane Mechanics and Function

Suspended pore-spanning membranes provide a promising route toward high-throughput biomimetic systems in which both sides of the bilayer remain accessible while membrane composition, asymmetry and geometry can be independently controlled (Figure 7). An important future direction is the development of single-molecule studies that directly test how membrane asymmetry and curvature regulate membrane-protein behaviour. Lipid asymmetry can be generated by assembling independently prepared monolayers, lipid-exchange approaches, or hemifusion-based methods^120,121^. Watanabe and colleagues demonstrated the parallel formation of more than 10,000 asymmetric bilayers with independently controlled leaflet compositions^121^. These strategies could be adapted to pore-spanning membranes using lipid exchange or GUV-bilayer hemifusion (**Figure 7A**). Such asymmetric suspended membranes could then be combined with single-molecule imaging to directly resolve lipid-protein reciprocal coupling (**Figure 7B**). The recently developed freestanding bilayer microscope enables unrestricted diffusion and single-particle tracking of integral membrane proteins and has been used to examine ion-channel open probability and protein dynamics in phase-separated membranes ^122^. Extending this approach to membranes with controlled asymmetry, composition, and curvature could enable single-molecule tracking or FRET measurements of protein diffusion, oligomerization, translocation, and conformational dynamics.

**Figure 7.**
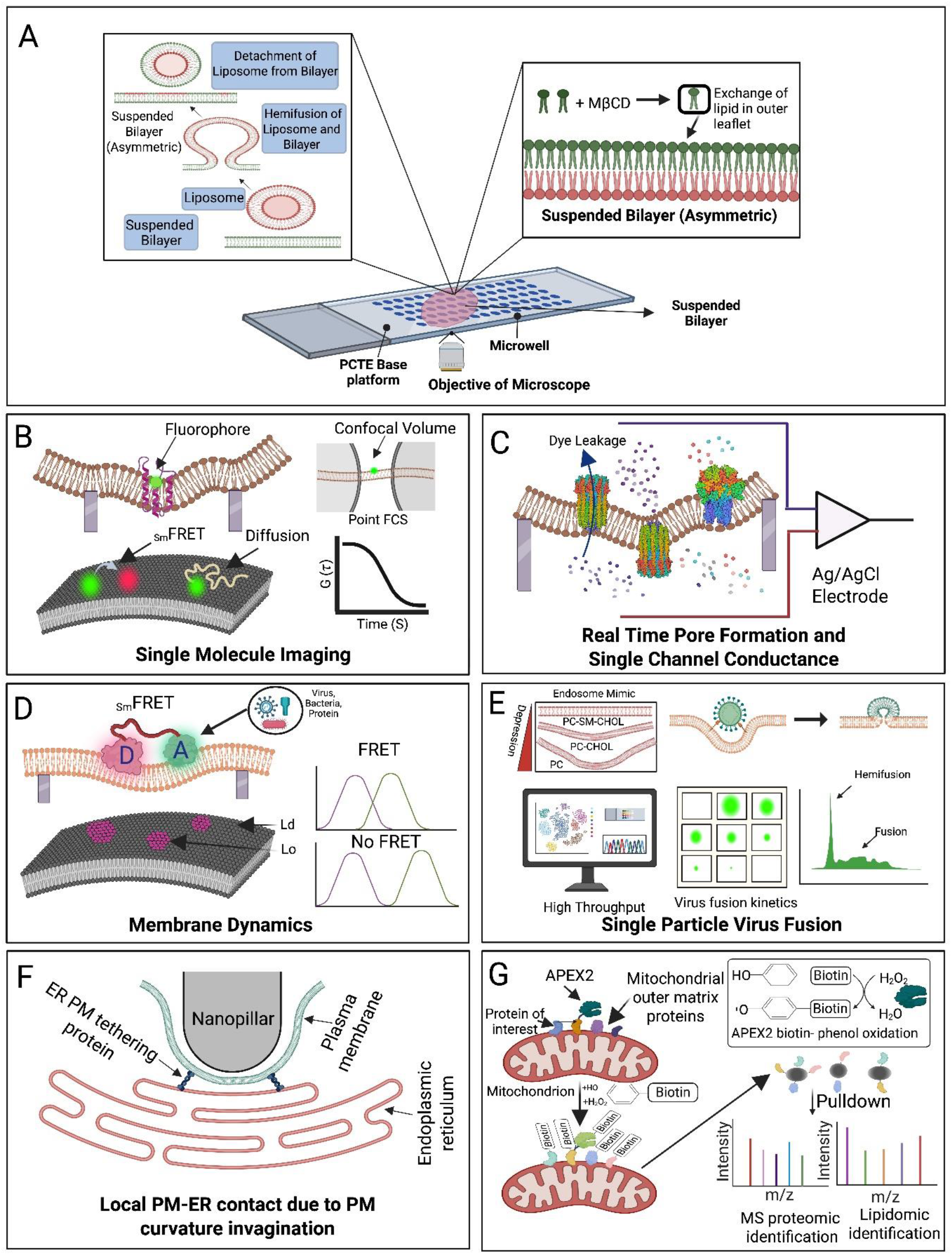
Experimental platforms to resolve lipid-protein coupling across membrane organization, dynamics and cellular architecture. **(A)** Engineering asymmetric membrane bilayers. A suspended bilayer can be generated on a porous polycarbonate track-etched (PCTE) membrane using liposome-bilayer hemifusion, followed by detachment of the liposome to generate a free-standing membrane. Controlled lipid asymmetry can subsequently be introduced by methyl-β-cyclodextrin (mβCD)-mediated lipid exchange into the outer leaflet, providing a platform to systematically control lipid composition, leaflet asymmetry and membrane organization. **(B)** Single-molecule imaging of protein-lipid interactions and membrane dynamics. Fluorescently labelled membrane proteins or lipids can be monitored at the single-molecule level to resolve molecular interactions, conformational changes and lateral diffusion. Single-molecule fluorescence resonance energy transfer (smFRET) reports protein conformational changes or intermolecular proximity, whereas fluorescence correlation spectroscopy (FCS) measures molecular diffusion and mobility. **(C)** Real-time pore formation and single-channel conductance. Suspended membranes containing pore-forming proteins can be combined with fluorescence-based dye leakage and Ag/AgCl electrode measurements to simultaneously monitor membrane permeabilization and quantify single-channel conductance, enabling analysis of pore formation and functional channel opening. **(D)** Membrane dynamics and lipid-domain-dependent protein behaviour. Phase-separated membranes containing liquid-disordered (L_d) and liquid-ordered (L_o) domains provide a controlled system for examining how membrane heterogeneity influences protein organization, interactions and conformational dynamics using smFRET. **(E)** Single-particle analysis of membrane fusion. Reconstituted membranes with defined lipid compositions, including PC-, PC-cholesterol and PC-SM-cholesterol membranes, can be used to study individual virus or virus-like particle fusion events. Single-particle fluorescence imaging and kinetic analysis can resolve fusion intermediates, including docking, hemifusion and fusion, and determine how lipid composition regulates fusion probability and kinetics. **(F)** Engineering membrane curvature to interrogate plasma membrane-ER coupling. Nanopillar-based platforms can impose defined nanoscale curvature on the plasma membrane, bringing the plasma membrane into close proximity with the endoplasmic reticulum (ER) and generating localized PM-ER contact sites. This provides a means to investigate how membrane curvature, lipid organization and ER-PM tethering proteins cooperate to regulate membrane contact-site formation. **(G)** Spatially resolved proteomics and lipidomics. APEX2-based proximity labelling can identify proteins associated with defined membrane microenvironments, followed by mass-spectrometric analysis. Coupling spatial proteomics with lipidomic profiling enables complementary mapping of proteins and lipids within specialized membrane regions, providing a framework to determine how local lipid composition and membrane architecture influence protein localization and function in living cells.

A second opportunity is the single-molecule measurement of transporter and ion-channel activity. Bilateral access to pore-spanning membranes allows ionic composition, pH, voltage, and ligands to be controlled independently on each side. Arrayed lipid bilayer chambers have already resolved quantized transport by individual α-hemolysin pores and ATP synthase molecules^123^, while membrane-on-a-chip devices have measured hundreds of individual channels and revealed heterogeneity in pore assembly and conductance ^124^. Incorporating controlled leaflet asymmetry and curvature into these platforms, as proposed in **Figure 7C**, could directly test how membrane architecture regulates channel opening, transporter activity, ion selectivity and drug response.

A third direction is the composition-dependent study of membrane phase behaviour together with membrane-protein function (**Figure 7D**). Suspended bilayers allow cholesterol, sphingomyelin, phospholipid saturation, and other lipid components to be varied systematically. In our previous work, we showed that sphingomyelin strongly enhances Cytolysin A (ClyA) pore-forming activity in suspended bilayers ^125^, while freestanding membrane arrays have shown that membrane-bound protein condensates reorganize lipids and reduce their mobility across opposing leaflets ^126^. High-throughput pore-spanning arrays could therefore map how lipid phase separation, asymmetry, and curvature jointly regulate protein partitioning, clustering, and activity.

The same platform would be particularly useful for studying pore-forming toxins. Suspended membranes have already enabled real-time analysis of ClyA and α-hemolysin pore formation ^57,124,125^. Importantly, the role of membrane curvature in regulating ClyA lipid binding and pore formation remains poorly explored. Arrays with different pore geometries could therefore provide controlled curvature while maintaining independent access to both membrane surfaces, allowing pore nucleation, oligomerization, conductance, and lipid scrambling to be followed in real time (**Figure 7C**).

Finally, freestanding endosomal membrane mimetics represent a promising platform for single-particle viral-fusion studies and antiviral discovery (**Figure 7E**). Suspended membrane chips have directly resolved H3N2 influenza fusion^124^, and suspended lipid bilayers have reproduced rapid dengue-virus fusion^125^. Future systems that reproduce endosomal lipid composition, leaflet asymmetry, cholesterol content, acidic pH, and membrane curvature could distinguish viral docking, hemifusion, pore opening, and content release at single-particle resolution. Integrating such measurements into pore arrays would enable parallel screening of fusion inhibitors. Overall, combining lipid composition, asymmetry, curvature, membrane-protein reconstitution, and single-molecule optical or electrical measurements in one high-throughput suspended platform could provide a next-generation approach for studying membrane mechanics, protein function, toxins, and viral entry. Together, these emerging platforms provide a route toward experimentally testing how lipid composition, asymmetry, curvature and mechanics jointly regulate protein organization and function.

### Nanopillar platforms: engineering membrane curvature to interrogate lipid-protein coupling in cell

An important future direction is to develop nanopillar-based membrane platforms that combine controlled nanoscale curvature with defined lipid compositions to quantitatively test lipid-protein interactions in physiologically relevant geometries (**Figure 7F**). Nanopillar and nanobar architectures have already emerged as powerful tools for imposing spatially defined membrane curvature in living cells and reconstituted membrane systems. Earlier studies demonstrated that cellular membrane deformation around nanopillars can selectively recruit curvature-sensitive proteins, including BAR-domain proteins and components of the clathrin-mediated endocytic machinery, establishing a direct connection between nanoscale membrane geometry and protein localization^127 106,128^. More recently, nanopillar-supported lipid-bilayer systems and NanoCurveS-type platforms have enabled quantitative measurements of protein curvature sensing, including the ability of F-BAR proteins such as FBP17 to discriminate membrane curvature at submicrometre length scales ^129,130^. Extending this concept to organelle organization, Yang *et al.* used vertical quartz nanopillars and nanobars to generate controlled plasma-membrane invaginations and showed that membrane curvature promotes the formation of ER-plasma-membrane contacts in cardiomyocytes. Their work demonstrated preferential recruitment of junctophilin-2 to curved membrane regions and identified Eps15 homology domain-containing proteins as curvature-sensing components that facilitate this targeting^131^. Thus, nanopillar systems provide an attractive bridge between reductionist membrane biophysics and cellular membrane organization. Future integration of nanopillar arrays with tunable lipid compositions, lipid-sensitive probes, super-resolution microscopy, single-molecule imaging and reconstituted proteins could allow systematic dissection of how lipid composition, membrane curvature and protein conformation cooperate to control membrane binding, oligomerization and function. Such platforms could ultimately transform nanopillars from tools for studying cellular curvature into quantitative, geometry-controlled assays for resolving the molecular principles of lipid-protein coupling across disease-relevant and physiological membrane environments.

### Spatial proteomics and lipidomics: mapping protein-lipid interactions in situ

A complementary frontier is the integration of spatial proteomics with spatial lipidomics to determine where proteins encounter specific lipid environments within cells (**Figure 7G**). Conventional proteomics and lipidomics largely require cellular extraction and therefore lose the spatial relationships that define membrane organization, whereas emerging mass-spectrometry imaging approaches can map hundreds of lipid species while retaining their cellular and subcellular distribution ^132–134^. In parallel, proximity-labeling strategies such as APEX2 and TurboID provide spatially restricted proteomic maps in living cells and have been successfully applied to membrane-bound organelles, membrane contact sites and lipid droplets^135–137^. Notably, APEX2-based proximity proteomics has enabled the identification of lipid-droplet-associated proteins while minimizing contamination from co-purifying organelles^137^. Combining these approaches with spatially resolved lipid measurements and lipid-protein crosslinking or proximity chemistry could link the identity and abundance of proteins to the lipid composition of their immediate membrane environment. Such multimodal strategies could distinguish proteins that merely occupy the same membrane from those whose localization is coupled to particular lipid species, while simultaneously revealing how local lipid composition varies across organelles, membrane contact sites and regions of distinct curvature. Recent advances in spatial multi-omics and high-resolution imaging mass spectrometry further suggest that simultaneous spatial mapping of proteins, lipids and metabolites is becoming increasingly feasible^133,138^. Integrating spatial proteomics and lipidomics with controlled-curvature nanopillar platforms could therefore provide a framework for determining where a protein encounters a lipid, which lipid species are present at that site, and how membrane geometry modulates their interaction, thereby connecting molecular recognition to membrane architecture and cellular function. The endoplasmic reticulum (ER), for example, is organized into morphologically and functionally distinct sheets, tubules and membrane contact sites, each associated with distinct molecular environments and cellular functions^134^. Recent spatial proteomics analysis of ER subdomains identified CLMN as an ER-tubule-enriched protein that links ER membranes to cortical F-actin at focal adhesions, illustrating how membrane morphology can be coupled to cytoskeletal organization, ER-plasma-membrane contacts and cell migration ^135^. Together, these approaches illustrate how membrane architecture can be resolved across multiple scales, from local lipid-protein interactions to organelle morphology and membrane-cytoskeletal interfaces.

## Conclusion

Membrane architecture provides an active physical context for protein organization and function. Lipid composition, transbilayer asymmetry, cholesterol, membrane curvature, thickness and mechanics collectively influence protein-membrane interactions through changes in packing, hydrophobic matching and local membrane geometry. Our quantitative analysis of 44 membrane proteins revealed substantial heterogeneity in curved-to-flat membrane association, with particularly strong curvature preference among several proteins known to sense or generate membrane curvature. In parallel, our hydrophobic-run analysis showed that disease-associated variants are enriched in shorter hydrophobic runs and that hydrophobic-to-polar substitutions occur frequently within transmembrane regions, linking protein sequence organization to hydrophobic matching and membrane compatibility. Together, these findings support lipid-protein reciprocal coupling as a framework in which membrane architecture regulates protein organization while proteins reciprocally reshape the local lipid environment.

These relationships can now be tested more directly using emerging experimental platforms that combine controlled lipid composition, leaflet asymmetry and membrane geometry with single-molecule imaging, lipid sensors, spatial proteomics and functional measurements. Suspended and pore-spanning membranes, nanopillar-based curvature platforms and spatially resolved lipidomic-proteomic approaches provide complementary routes for separating the individual and combined contributions of lipid composition, curvature, asymmetry and mechanics. Integrating these approaches with quantitative analyses of protein curvature preference and hydrophobic sequence organization should help define how membrane architecture encodes physical information that is translated into protein localization, conformation and activity, and how disruption of this coupling contributes to disease-associated membrane dysfunction.

## Materials and Methods

### Computational methods

The quantitative analyses presented in this study were performed using publicly available protein structural and sequence databases. Two complementary computational analyses were carried out. First, membrane-protein orientation and membrane association were evaluated using the Orientation of Proteins in Membranes (OPM) framework and the PPM 3.0 web server. This analysis was used to compare the apparent membrane association of structurally characterized proteins in planar and curved membrane environments. Second, hydrophobic and hydrophilic residue runs were quantified from protein sequences obtained from the SCOP (Structural Classification of Proteins), OPM, and MutHTP databases.

### Selection and analysis of membrane-protein structures

A structurally diverse set of 44 membrane proteins (Table 1) was selected for quantitative analysis of membrane association and curvature preference. Protein structures were selected based on the availability of experimentally determined structural coordinates and membrane-orientation information. The analyzed set included representative membrane proteins spanning different structural classes, membrane topologies, and functional categories. The complete list of proteins, corresponding PDB identifiers, and membrane-association values used for the analysis is provided in Table 1.

Structural coordinates and membrane-positioning calculations were obtained using the OPM PPM 3.0 web server (https://opm.phar.umich.edu/ppm_server3). PPM 3.0 permits positioning of protein structures in planar or curved membrane geometries and determines energetically optimal membrane arrangements and associated transfer-energy parameters. The calculations were performed using the PPM 3.0 version of the server.

To gain insight into how diverse membrane proteins interact with the membrane, we used a computational membrane-positioning approach. Protein orientations in membranes were theoretically calculated by minimizing the transfer energy of a protein from water to a planar slab that serves as an approximation of the membrane hydrocarbon core. The membrane binding propensity was calculated by submitting the coordinate information of each protein structure to the OPM/PPM 3.0 server. In these calculations, a protein was treated as a rigid body that freely floats within the planar hydrocarbon core of a lipid bilayer. The accessible surface area is calculated using the SOLVA subroutine of NACCESS, with Chothia atomic radii and without hydrogen atoms. OPM uses solvation parameters derived specifically for lipid bilayers and normalized by the effective concentration of water, which changes gradually along the bilayer normal within a relatively narrow region between the lipid headgroup regions and the hydrocarbon core.

For proteins for which multiple experimentally determined structures or conformational states were available, individual structures were treated as separate structural observations when they represented distinct membrane-associated conformations analyzed in the manuscript. Structures lacking the coordinate information required for the analysis were excluded.

### PPM 3.0-based membrane positioning and apparent membrane association

The spatial arrangement of each selected protein structure relative to the membrane was calculated using PPM 3.0. Protein coordinate files were provided in PDB format, and the appropriate membrane and topology settings were selected according to the membrane-associated state being evaluated. PPM 3.0 determines the energetically optimal spatial position of a protein relative to a planar or curved membrane by minimizing the protein transfer energy in the membrane environment. The server can explicitly allow membrane curvature and determine whether a protein structure is better accommodated by a planar or curved membrane geometry.

For each protein structure, the apparent membrane-association or water-to-membrane transfer-energy value obtained from PPM 3.0 was recorded for the corresponding planar and curved membrane calculations. The values reported in Table 1 were used for subsequent quantitative comparison. PPM 3.0 calculations were performed using the same calculation settings for the structures included in the comparative analysis, except where topology or membrane assignment required structure-specific settings.

The PPM 3.0 framework accounts for the energetic contributions associated with positioning protein atoms within the anisotropic membrane environment and provides transfer-energy and membrane-positioning parameters. The present analysis uses these calculated energetic values as a relative measure of membrane association and curvature preference rather than as an experimental measurement of absolute membrane-binding affinity.

### Quantification of curvature preference

To quantify relative preference for curved versus planar membrane environments, the apparent membrane-association values obtained for the two membrane geometries were compared for each protein. The curvature-enrichment ratio was calculated as the ratio of the magnitude of the calculated transfer-energy values for the curved and planar membrane states:

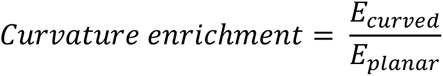

where *E_curved_* and *E_planar_* represent the PPM 3.0-derived apparent membrane-association/transfer-energy values for the curved and planar membrane states, respectively.

The corresponding log2 fold-enrichment was calculated as:

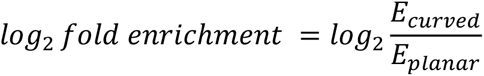

*log_2_ ^fold enrichment^* greater than zero therefore indicates a higher magnitude of calculated membrane-transfer energy in the curved state, whereas a value below zero indicates greater apparent association with the planar state. A value of zero corresponds to equivalent calculated association between the two membrane geometries.

The resulting values were used to generate the quantitative curvature-association analysis shown in the manuscript. Because these values are derived from a computational membrane-positioning model, they are interpreted as relative curvature preference based on the calculated transfer energies and not as direct experimental measurements of membrane-binding affinity.

### Protein structural and functional annotation

Protein names, structural classifications, PDB identifiers, and functional descriptions were obtained from the corresponding structural databases and the primary literature associated with each protein (Table 1). When multiple structures of the same protein were included, the specific PDB accession and structural state were retained to distinguish individual observations.

### Analysis of hydrophobic and hydrophilic residue runs

Protein sequences were obtained from the Structural Classification of Proteins (SCOP) database hosted by the Protein Data Bank in Europe (PDBe) (https://www.ebi.ac.uk/pdbe/scop/), the OPM database (https://opm.phar.umich.edu/), and the MutHTP database (https://www.iitm.ac.in/bioinfo/MutHTP/). The corresponding database resources were used as the source datasets for the sequence-based analyses described below.

Each protein sequence was converted into a binary representation according to the hydrophobic or hydrophilic character of its constituent amino acids. The hydrophobic residue set was defined as Ala (A), Phe (F), Gly (G), Ile (I), Leu (L), Met (M), Pro (P), Val (V), and Trp (W), whereas the hydrophilic residue set was defined as Cys (C), Asp (D), Glu (E), His (H), Lys (K), Asn (N), Gln (Q), Arg (R), Ser (S), Thr (T), and Tyr (Y). Hydrophobic residues were represented by 1 and hydrophilic residues by 0.

The lengths of consecutive runs of 0 and 1 were determined for each binary-encoded protein sequence. Thus, each continuous sequence of hydrophobic residues was recorded as a hydrophobic run, whereas each continuous sequence of hydrophilic residues was recorded as a hydrophilic run. The distribution of run lengths was subsequently determined for each protein and aggregated according to the corresponding protein class or database category.

The run analysis was used as a sequence-level measure of the organization and prevalence of consecutive hydrophobic or hydrophilic residues. A hydrophobic run was not assumed to represent a transmembrane helix solely on the basis of its sequence length.

### SCOP dataset analysis

Protein sequences classified in the SCOP database were analyzed to evaluate the distribution of hydrophobic and hydrophilic runs among structurally classified proteins. The SCOP database available through PDBe was used as the source of structural classification and sequence information.

Protein sequences were processed using the binary residue-encoding and run-length procedure described above. The analyzed protein categories and the number of sequences included in each category are collected from SCOP database membrane family proteins. Where duplicate entries, incomplete sequences, or redundant records were excluded, the corresponding filtering criteria were applied consistently across the dataset. Run lengths were calculated independently for each protein before aggregation across the corresponding SCOP category so that the analysis retained protein-level information.

### OPM sequence dataset analysis

Protein sequences associated with the OPM database were analyzed using the same binary sequence-encoding and run-length procedure. Protein entries were assigned to the corresponding OPM categories based on the database annotations available at the time of analysis. The structural details of the transmembrane proteins included in each category are provided in Table 1. Hydrophobic and hydrophilic run lengths were calculated independently for each sequence, and the resulting distributions were compared among the analyzed protein categories.

### MutHTP disease-associated mutation analysis

Disease-associated protein mutation information was obtained from the MutHTP database (Mutations in Human Transmembrane Proteins; https://www.iitm.ac.in/bioinfo/MutHTP/). MutHTP contains disease and neutral mutation information for human transmembrane proteins and includes missense, deletion, and insertion mutations. For the present analysis, missense mutations were analyzed. For each analyzed protein, the reported mutation position was mapped onto the corresponding protein sequence. A mutation was classified as occurring within a hydrophobic or hydrophilic run when the affected residue was located within a consecutive run identified by the sequence-based analysis described above. For substitution analysis, amino acids were grouped into hydrophobic and hydrophilic classes using the same residue definitions described above. Substitutions were categorized as hydrophobic-to-hydrophobic, hydrophilic-to-hydrophilic, hydrophobic-to-hydrophilic, or hydrophilic-to-hydrophobic. The frequencies of these substitution classes were calculated relative to the total number of substitutions analyzed.

### Analysis of long hydrophobic runs

For analyses in which long hydrophobic segments were specifically examined, hydrophobic runs were classified according to their consecutive-residue length. The threshold used in the manuscript was greater than 16 consecutive hydrophobic residues. This threshold was applied consistently across the analyzed datasets. For each protein category, the frequency of proteins containing hydrophobic runs exceeding this threshold was determined and compared with the corresponding distributions in other protein groups.

### Data analysis and visualization

The sequence-based analyses consisted of sequence retrieval, residue classification, binary encoding, run-length identification, dataset categorization, quantitative aggregation, and visualization. For the curvature analysis, each analyzed structural entry was treated as an individual observation. The calculated curvature-enrichment and log2 fold-enrichment values were used for visualization and quantitative comparison. The number of structures/proteins included in each analysis is reported in the corresponding figure legends.

### Reproducibility and data availability

The complete list of protein identifiers, PDB accession numbers, PPM 3.0-derived membrane-association values, curvature-enrichment ratios, and log2 fold-enrichment values used for the curvature analysis are provided in Table 1 and figure legends. All structural and sequence information analyzed in this study was obtained from publicly accessible resources, including the OPM/PPM 3.0 server, SCOP, and MutHTP. The computational procedures described above were applied consistently to the corresponding datasets.

### Experimental Methods

#### Suspended lipid bilayer preparation

Pore-spanning suspended lipid bilayers (SuLBs) were prepared using hydrophilic polycarbonate track-etched (PCTE) membrane substrates. PVP-coated hydrophilic PCTE membranes were placed on thoroughly cleaned glass coverslips. Giant unilamellar vesicles (GUVs) of the required lipid composition were then added to the PCTE surface and incubated at 37 °C for 30 min to promote vesicle rupture and spreading of the lipid bilayer across the pores. Excess lipid material was removed by gentle washing with buffer. Formation of the pore-spanning bilayers was verified by fluorescence microscopy and, where required, by atomic force microscopy (AFM) or total internal reflection fluorescence (TIRF) microscopy.

For fluorescence-based membrane-permeabilization measurements, sulforhodamine B (SRB) was confined within the PCTE microwells before membrane sealing. A 1-µm PCTE membrane was placed on a cleaned glass coverslip, and SRB solution was introduced into the membrane pores. A GUV suspension was then added together with CaCl₂ to promote vesicle rupture and bilayer spreading over the pores. The membrane assembly was incubated at 37 °C for 30 min and gently washed before imaging. Following addition of the membrane-active protein, loss of SRB fluorescence from individual membrane-sealed microwells was monitored over time as a measure of membrane permeabilization.

#### Generation of membrane asymmetry by mβCD-mediated lipid exchange and hemifusion

The asymmetric-membrane strategy illustrated in Figure 7A was constructed from previously established approaches for generating transbilayer lipid asymmetry. These methods were not performed experimentally in the present study but were used as the basis for the proposed asymmetric suspended-membrane platform.

One established approach uses methyl-β-cyclodextrin (mβCD)-mediated lipid exchange. Visco, Chiantia and Schwille developed a method in which supported lipid bilayers formed by vesicle fusion were exposed to lipid-loaded mβCD complexes. In this system, mβCD acts as a lipid carrier and enables selective enrichment of the solvent-exposed upper leaflet while largely preserving the composition of the leaflet facing the solid support. Using sphingomyelin as the exchanged lipid, the asymmetric supported lipid bilayers were generated and confirmed leaflet-specific differences in lipid mobility by fluorescence correlation spectroscopy. The mβCD-mediated lipid-exchange concept shown schematically in Figure 7A was therefore adapted from the method described by Visco et al^139^.

A complementary approach uses controlled hemifusion to generate asymmetric giant unilamellar vesicles. Enoki and Feigenson developed a method in which a GUV of one lipid composition is brought into contact with a supported lipid bilayer of a different composition. Calcium promotes hemifusion between the two contacting membranes, allowing lipid diffusion and exchange between the fused outer leaflets without complete fusion of the bilayers. After separation from the supported membrane, the resulting GUV retains a different lipid composition in its outer and inner leaflets. Using leaflet-specific fluorescent probes, the authors showed that nearly all newly exchanged lipids were localized to the outer leaflet and that high levels of lipid exchange could be achieved under optimized conditions. The hemifusion-based asymmetry concept represented in Figure 7A was therefore adapted from the method reported by Enoki and Feigenson^140^.

#### Nanopillar-based membrane-curvature platform

The nanopillar configuration illustrated in Figure 7F was developed as a conceptual extension of established membrane-curvature platforms, including the system reported by Yang *et al*^131^. In that study, vertical quartz nanopillars were used to impose defined nanoscale curvature on the plasma membrane of cultured cells, generating highly curved inward membrane invaginations that could be compared with neighboring, relatively flat membrane regions.

Fluorescence imaging together with structural imaging, including focused ion beam-scanning electron microscopy, was used to examine how membrane curvature influences plasma-membrane–endoplasmic-reticulum contact formation and the recruitment of curvature-responsive proteins such as junctophilin-2 and Eps15 homology domain-containing proteins^131^. Building on this established experimental principle, Figure 7F presents a proposed framework for extending nanopillar-based curvature control to studies of lipid–protein reciprocal coupling. In this design, defined membrane geometry could be combined with controlled lipid composition, lipid-sensitive probes, and measurements of protein localization or organization to examine how membrane curvature and the local lipid environment jointly regulate protein behavior.

#### Literature-Derived Data Collection and Analysis

Figures 1 and 2 were prepared by combining published quantitative information with schematic representations developed for the present study. Figure 1 summarizes established concepts in membrane evolution and environmental adaptation. The homeoviscous-adaptation scheme in Figure 1C was based on the classic work of Sinensky, which showed temperature-dependent changes in fatty-acid saturation and chain length that help maintain membrane viscosity^19^. Figure 1D summarizes the pressure-dependent homeocurvature response reported in deep-sea invertebrates by Winnikoff *et al*^20^. For Figure 2A–D, leaflet-specific lipid composition, phospholipid unsaturation, and abundant lipid species were reconstructed from the published quantitative erythrocyte plasma-membrane lipidomic data of Lorent *et al*^27^. Figure 2E was drawn as a schematic representation of lipid molecular geometry and preferred membrane curvature, whereas Figure 2F was constructed to illustrate the relationship between leaflet-specific lipid packing and transmembrane-domain architecture described in the same study ^27^. Figures 5–7 contain schematic representations of established membrane-analysis approaches and published membrane-organizing principles that were integrated into the framework developed in the present manuscript. Figure 5 compares commonly used biomimetic membrane platforms discussed in the corresponding section of the manuscript. Figure 6 summarizes established lipid-sensing strategies and cholesterol organization; in particular, the accessible, sphingomyelin-sequestered, and residual cholesterol pools represented in Figure 6B–C were based on the cholesterol-accessibility framework reported by Das *et al*^107^. Figure 7B–E and 7G were constructed as proposed extensions of established single-molecule imaging, membrane-conductance, phase-behavior, membrane-fusion, and spatial proteomic/lipidomic approaches described and cited in the corresponding sections of the manuscript. These panels were redrawn for the present study to integrate the relevant published methods and concepts within the broader lipid–protein reciprocal-coupling framework.

## Abbreviations

AFM: Atomic force microscopy
APEX2: Engineered ascorbate peroxidase 2
BAR: Bin/Amphiphysin/Rvs domain
ClyA: Cytolysin A
DAG: Diacylglycerol
DHAP: Dihydroxyacetone phosphate
ER: Endoplasmic reticulum
F-BAR: Fes/CIP4 homology-BAR domain
FCS: Fluorescence correlation spectroscopy
FRET: Förster resonance energy transfer
FTIR: Fourier-transform infrared spectroscopy
G1P: sn-Glycerol-1-phosphate
G3P: sn-Glycerol-3-phosphate
GPMV: Giant plasma membrane vesicle
GUV: Giant unilamellar vesicle
Ld: Liquid-disordered phase
Lo: Liquid-ordered phase
LUV: Large unilamellar vesicle
mβCD: Methyl-β-cyclodextrin
N-BAR: N-terminal amphipathic helix-containing BAR domain
PA: Phosphatidic acid
PC: Phosphatidylcholine
PCTE: Polycarbonate track-etched
PE: Phosphatidylethanolamine
PH: Pleckstrin homology domain
PI: Phosphatidylinositol
PI(4,5)P₂ / PIP₂: Phosphatidylinositol 4,5-bisphosphate
PM: Plasma membrane
PS: Phosphatidylserine
SLB: Supported lipid bilayer
SM: Sphingomyelin
SM-CHOL: Sphingomyelin-cholesterol
smFRET: Single-molecule Förster resonance energy transfer
SUV: Small unilamellar vesicle
TMD: Transmembrane domain

## Author Contributions

A.S. conceived and led the project. A.S. and M.S.A.M. performed the quantitative and computational analyses. M.S.A.M., D.S., N.D., D.B. and A.S. contributed to figure preparation, interpretation of the results, and writing and revision of the manuscript. All authors read and approved the final manuscript.

## Acknowledgements

A.S. acknowledges the Anusandhan National Research Foundation (ANRF), Government of India, for support through the National Postdoctoral Fellowship (PDF/2020/000678).

